# Reinforcement regulates timing variability in thalamus

**DOI:** 10.1101/583328

**Authors:** Jing Wang, Eghbal Hosseini, Nicolas Meirhaeghe, Adam Akkad, Mehrdad Jazayeri

## Abstract

Learning reduces variability but variability can facilitate learning. This paradoxical relationship has made it challenging to tease apart sources of variability that degrade performance from those that improve it. We tackled this question in a context-dependent timing task requiring humans and monkeys to flexibly produce different time intervals with different effectors. Subjects’ timing variability featured two novel and context-specific sources of variability: (1) slow memory-contingent fluctuations of the mean that degraded performance, and (2) fast reinforcement-dependent regulation of variance that improved performance. Signatures of these processes were evident across populations of neurons in multiple nodes of the cortico-basal ganglia circuits. However, only in a region of the thalamus involved in flexible control of timing were the slow performance-degrading fluctuations aligned to performance-optimizing regulation of variance. These findings provide direct evidence that the nervous system makes strategic use of exploratory variance to guard against other undesirable sources of variability.

## Introduction

While interacting with a dynamic and uncertain environment, humans and animals routinely rely on learning through trial and error to optimize their behavior. This type of learning is most rigorously formulated within the framework of reinforcement learning (RL). The key idea behind RL is to compute a value function for all possible actions based on previous outcomes, and use those action-outcome relationships to guide future actions (Sutton and Barto, 1998). RL has been extremely valuable in explaining behavior when an agent has to choose among a small set of discrete options, such as classic multi-armed bandit problems (Daw et al., 2005). However, learning through trial and error is also critical for other behavioral settings where the action space is large and continuous (Dhawale et al., 2017). For example, this strategy seems particularly relevant when the agent has to adjust its movements based on a binary or scalar (unsigned) feedback (Chen et al., 2017; Dam et al., 2013; Izawa and Shadmehr, 2011; Nikooyan and Ahmed, 2015; Pekny et al., 2015; Shmuelof et al., 2012a; Wu et al., 2014). In this case, classic RL is not feasible as it requires the agent to compute the value function over an infinity of movement parameters (e.g., all the kinematic degrees of freedom associated with a simple reach and grasp movement). Moreover, in a continuous state space, the slightest source of variability such as the inherent noise in the motor system would degrade the agent’s ability to update action-outcome relationships.

This raises the question of how humans and animals employ trial and error learning when the space of possibilities is continuous. One intriguing hypothesis is that the brain directly regulates variability to facilitate learning (Huang et al., 2011; Kao et al., 2005; Olveczky et al., 2005; Tumer and Brainard, 2007). The basic idea is to reduce variability when a trial yields reward (i.e., exploit previously rewarded action), and increase variability in the absence of reward (i.e., promote exploration of new possibilities). This strategy captures the spirit of RL in moderating exploration versus exploitation without explicitly computing a continuous value function. Several indirect lines of evidence support this hypothesis (Dhawale et al. 2019; Shmuelof et al. 2012; Vaswani et al. 2015; Cashaback et al. 2017; Palidis et al.; van der Kooij and Smeets 2019; Cashaback et al. 2019; Chen et al. 2017; Dam et al. 2013; Izawa and Shadmehr 2011; Nikooyan and Ahmed 2015; Pekny et al. 2015; Shmuelof et al. 2012; Wu et al. 2014). For example, humans learn more efficiently if the structure of their natural movement variability aligns with the underlying learning objective (Wu et al., 2014), reaching movements become more variable during periods of low success rate (Izawa and Shadmehr, 2011; Pekny et al., 2015), and metrics of saccadic eye movements become more variable in the absence of reward (Takikawa et al., 2002).

However, the relationship between variability and learning is nuanced. While increasing motor variability may play a direct role in the induction of learning (Dhawale et al., 2017), one of the primary functions of motor learning is to reduce variability (Crossman, 1959; Harris and Wolpert, 1998; Smith et al., 2006; Sternad and Abe, 2010; Thoroughman and Shadmehr, 2000; Verstynen and Sabes, 2011). This two-sided relationship between learning and variability has made it challenging to tease apart behavioral and neural signatures of variability that degrade performance from those that improve it. A rigorous assessment of the interaction between learning and variability demands two important developments. First, we need a method capable of teasing apart sources of variability that hinder performance from those that facilitate learning. Second, we need to verify that the underlying neural circuits rely on reinforcement to regulate the variability along task-relevant dimensions. To address these problems, we focused on a motor timing task in which monkeys produced different time intervals using different effectors. We first verified the presence of motor variability due to memory drift and reward-dependent adjustment in motor timing. We then developed a generative model to explain how reward regulates variability and facilitates learning. Finally, we probed the underlying neural circuits and asked whether reward could regulate task-relevant variability within the nervous system. We recorded from multiple nodes of the cortico-basal ganglia circuits that were previously found to support monkeys’ timing behavior (Wang et al., 2018). Results indicated that the variability across the population of thalamic neurons with projections to the dorsomedial frontal cortex (DMFC) was indeed regulated by reward along task-relevant dimensions in a context-specific manner.

## Results

### Cue-Set-Go task

Two monkeys were trained to perform a Cue-Set-Go (CSG) motor timing task (Figure 1A). On each trial, the animal aimed to produce either an 800 ms (Short) or a 1500 ms (Long) time interval (*t*_*t*_) either with a saccade (Eye) or with a button press (Hand). Each trial was initiated by presenting a fixation cue (“Cue”) at the center of the screen. The Cue consisted of a circle and a square. A combination of color and shape indicated the trial type. The four trial types, Eye-Short (ES), Eye-Long (EL), Hand-Short (HS), and Hand-Long (HL) were randomly interleaved throughout the session. After a random delay, a visual stimulus (“Tar”) was flashed to the left or right of the screen. This stimulus specified the position of the saccadic target for the Eye trials and served no function in the Hand trials. After another random delay, the presentation of a “Set” flash around the fixation spot indicated the start of the timing period. Animals had to proactively initiate a saccade or a button press aiming at the desired *t*_*t*_. We refer to the movement initiation as the “Go”. We measured the produced interval, *t*_*p*_, as the interval between Set and Go. To receive reward, the relative error, defined as *e* = (*t*_*p*_-*t*_*t*_)/*t*_*t*_, had to be within a window (Figure 1A). On rewarded trials, the magnitude of reward decreased linearly with the size of the error. The width of the reward window was controlled independently for each trial type and was adjusted adaptively on a trial-by-trial basis (see Methods).

**Figure 1.**
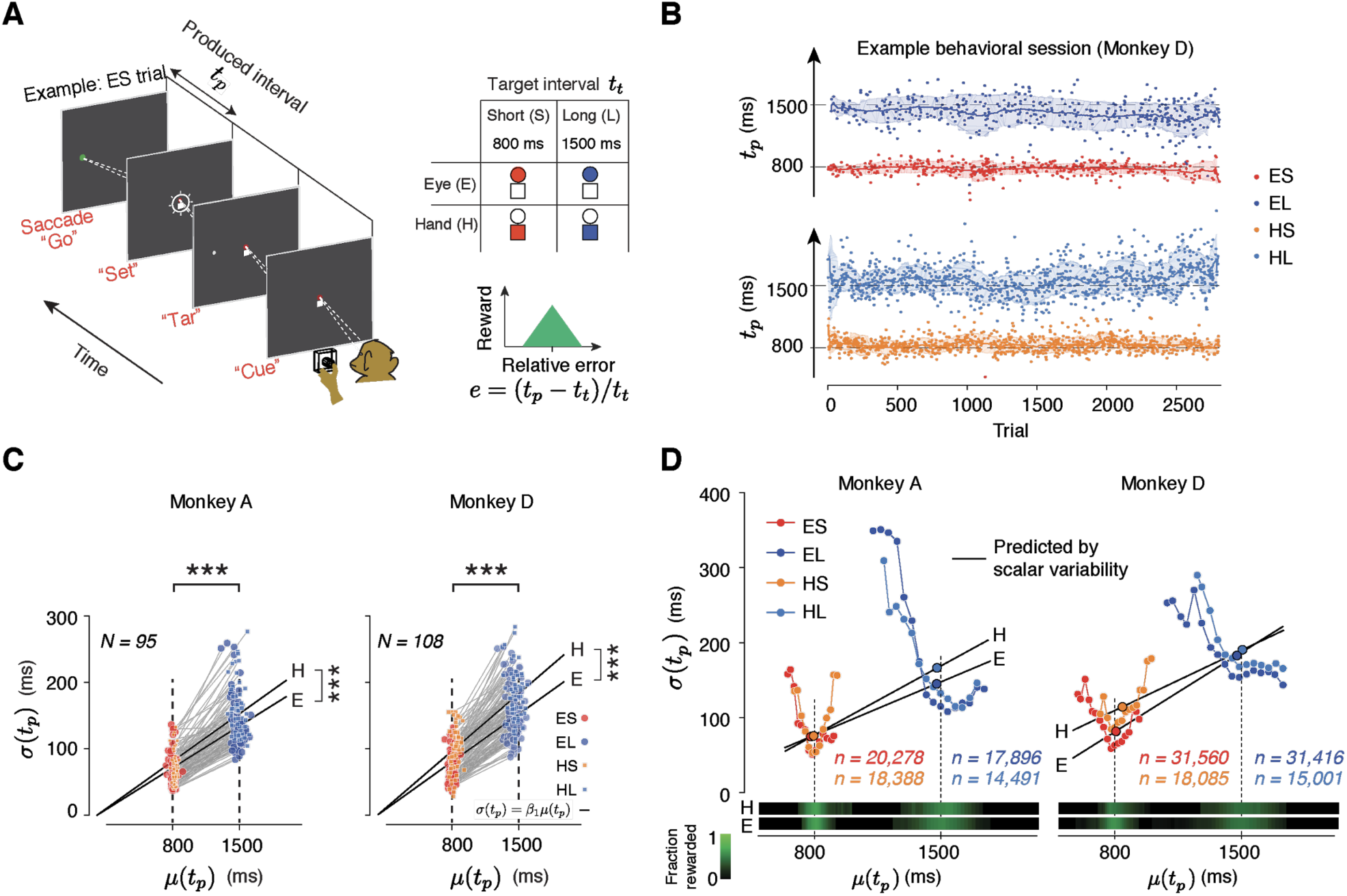
Task, behavior, and reward-dependency of variability. **(A)** The Cue-Set-Go task. **(B)** Animal’s behavior in a representative session. For visual clarity, *t*_*p*_ values (dots) for different trial types are shown along different abscissae for each trial type. The solid line and shaded area are the mean and standard deviation of *t*_*p*_ calculated from a 50-trial sliding window. **(C)** Standard deviation of *t*_*p*_ (*σ*(*t*_*p*_)) as a function of its mean (*μ*(*t*_*p*_)) for each trial type in each behavioral session. Each pair of connected dots corresponds to Short and Long of the same effector in a single session. In both animals, the variability was significantly larger for the Long compared to the Short for both effectors (one-tailed paired-sample t test, *** p <<0.001, for monkey A, n =190, t_128_ = 157.4; for monkey D, n = 216, t_163_ = 181.7). The solid black lines show the regression line relating *σ*(*t*_*p*_) to *μ*(*t*_*p*_) across all behavioral sessions for each trial type (*σ*(*t*_*p*_) = *β*_*1*_ *μ*(*t*_*p*_)). Regression slopes were positive and significantly different from zero for both effectors (*β* = 0.087 ± 0.02 mean ± std for Eye and 0.096 ± 0.021 for Hand in Monkey A; *β*_*1*_ = 0.10 ± 0.02 for Eye and 0.12 ± 0.021 for Hand in Monkey D). Hand trials were more variable than Eye ones (one-tailed paired-sample t-test, for monkey A, n = 95, t_52_ = 6.92, *** p <<0.001, and for monkey D, n = 108, t_61_ = 6.51, ***p << 0.001). **(D)** Dependence of timing variability on reward rate. Top: The plot shows the relationship between the standard deviation and mean of *t*_*p*_ (*σ*(*t*_*p*_) and *μ*(*t*_*p*_)) when these statistics are derived locally from the full distribution (Figure S1). The larger circles with black outline correspond to grand *σ*(*t*_*p*_) and *μ*(*t*_*p*_) computed from combining *t*_*p*_ values for each trial type across all behavioral sessions. The structure of variability (small circles connected with colored lines) was non-stationary, reward-dependent, and distinctly different from predictions of scalar variability (black line). Bottom: Relative reward rate is shown for each *μ*(*t*_*p*_) bin. The fraction rewarded was the ratio between the number of rewarded trials and total number of trials across all behavioral sessions

Animals learned to use the Cue and flexibly switched between the four trial types (Figure 1B). For both effectors, a robust feature of the behavior was that produced intervals (*t*_*p*_) were more variable for the Long compared to Short (Figure 1C). This is consistent with the common observation that timing variability scales with the interval being timed (Gibbon, 1977; Malapani and Fairhurst, 2002). This *scalar variability* was manifest in the linear relationship between mean (*μ*) and standard deviation (*σ*) of *t*_*p*_ in each behavioral session (Figure 1C, the slope of the linear test was significantly positive, one-tailed t-test, p << 0.001, for monkey A, df = 128, t = 32.12; for monkey D, p << 0.001, df = 163, t = 24.06).

### Deconstructing motor timing variability

The neurobiological basis of *scalar variability* is not understood. Models of interval timing have attributed scalar variability to a variety of processes including a variable clock, a noisy accumulation process, noisy oscillations, and errors related to storage and/or retrieval of an interval (Church and Broadbent, 1990; Gibbon et al., 1984; Grossberg and Schmajuk, 1989; Jazayeri and Shadlen, 2010; Killeen and Fetterman, 1988; Machado, 1997; Oprisan and Buhusi, 2014; Simen et al., 2011; Staddon and Higa, 1999). These models assume that variability is stationary and therefore, predict that *σ*(*t*_*p*_) has a fixed linear relationship to *μ*(*t*_*p*_). To test this assumption, we analyzed local estimates of *σ*(*t*_*p*_) as a function of *μ*(*t*_*p*_) across trials of the same type from blocks of 50 consecutive trials. The relationship between *μ*(*t*_*p*_) and *σ*(*t*_*p*_) across blocks of trials was strikingly different from the predictions of scalar variability (Figure 1D, S1): local estimates of *σ*(*t*_*p*_) decreased when *μ*(*t*_*p*_) was close to the desired *t*_*t*_, and increased when *μ*(*t*_*p*_) deviated from *t*_*t*_. In other words, variability was smaller when the animal received more reward and vice versa (Figure 1D). This inverse relationship was readily evident from the strong negative correlation between *σ*(*t*_*p*_) and the fraction of reward (p << 0.001, *r* = −0.48 for Monkey A and *r* = −0.50 for Monkey D). Although this result rejects models of interval timing that assume stationarity, the fact that animals’ behavior is modulated by reward is unsurprising. In our experiment, the only information animals could rely on to calibrate their behavior was the amount of reward. The relationship between reward and variability suggests that animals calibrate their timing behavior through ongoing adjustment of behavioral variability.

The non-stationarity of behavior was also evident from serial correlations of *t*_*p*_ between trials of the same type (e.g., all trials of ES within a session). In all four trial types, *t*_*p*_ exhibited significant serial correlations up to a trial lag of 20 or more (p < 0.01, dash lines: 1% and 99% confidence bounds by estimating the null distribution from shuffled series, Figure 2A). These fluctuations have been reported in many tasks (Gilden et al., 1995; Merrill and Bennett, 1956; Weiss et al., 1955) and can be relatively strong in movements (Chaisanguanthum et al., 2014), reaction times (Laming, 1979), and interval timing (Chen et al., 1997; Murakami et al., 2017). Importantly, these correlations were context-specific; i.e. serial correlations were stronger between trials with the same effector and same *t*_*t*_ compared to trials that were associated with different *t*_*t*_ and/or different effector (see Figure 2B for statistics and Figure S2 for serial correlations between all trial types and saccade directions). The context-specificity of these correlations suggests that they were not solely due to non-specific fluctuations of internal states such as the overall level of alertness, which should persist across the four randomly interleaved trial types.

**Figure 2.**
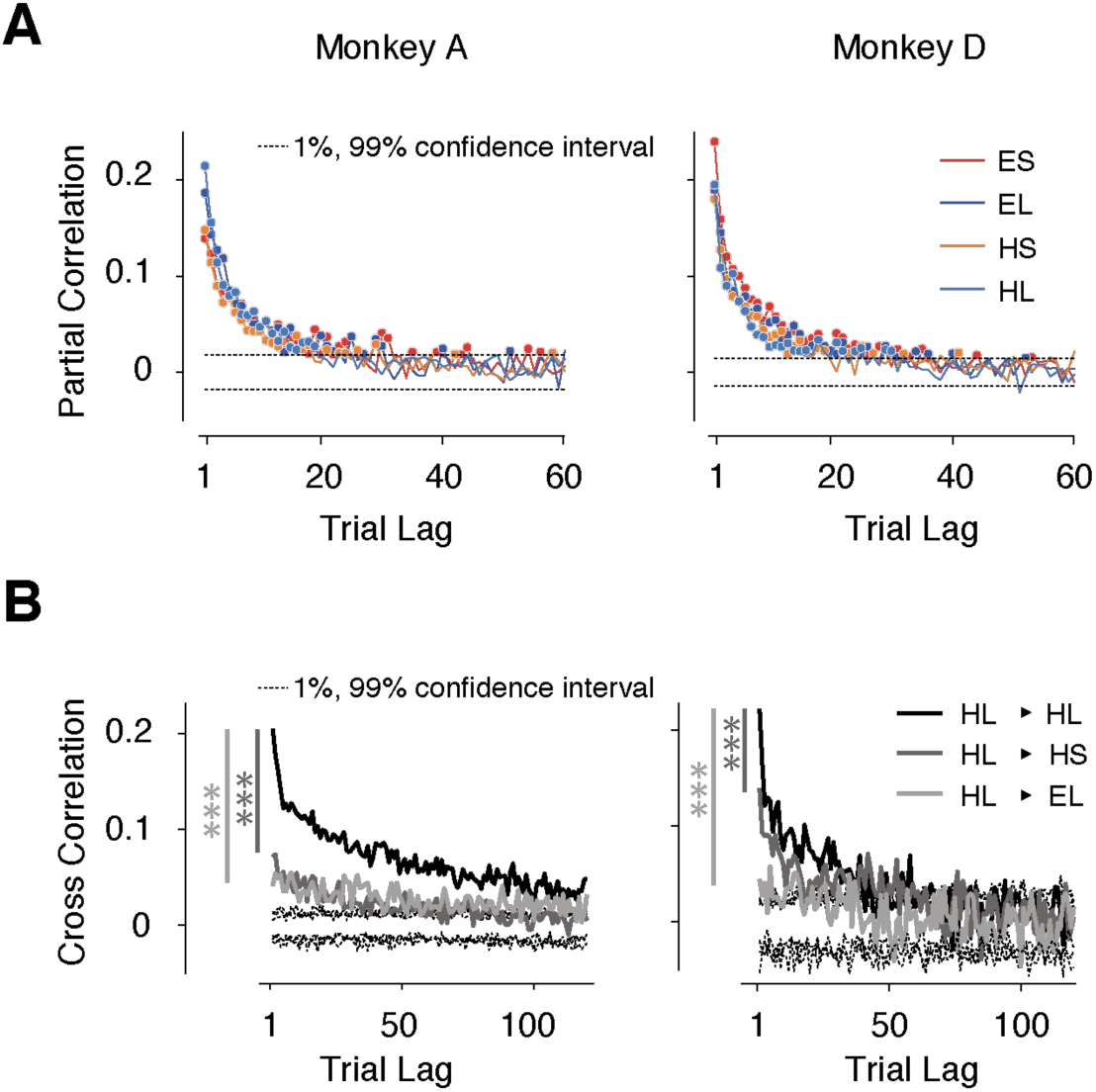
Context-dependent slow fluctuations of timing intervals. **(A)** Long-term correlation of *t*_*p*_ across trials of the same type (same effector and target interval). For each behavioral session, and each trial type, we computed the partial correlation of *t*_*p*_ as a function of trial lag. Each curve shows the average partial correlations across all behavioral sessions. Four types of trials are shown in different colors. Filled circles: correlation values that are larger than the 1% and 99% confidence interval (dashed line). **(B)** Examples showing the Pearson correlation coefficients of *t*_*p*_*’s* as a function of trial lag. HL-HL indicates the correlation was averaged across trials transitioning between HL and HL; 1% and 99% confidence intervals were estimated from the null distribution. Serial correlations were stronger between trials with the same effector and interval compared to trials with the same effector but different interval (*** dark gray, paired sample *t*-test on two sets of cross correlations with less than 20 trial lags and combining 4 trial types; monkey A: p << 0.001, n = 80, t_79_ = 9.8; monkey D: p << 0.001, n = 80, t_79_ = 5.8), and trials of the different effector but same interval (*** light gray, monkey A: p << 0.001, n = 80, t_79_ = 6.7; monkey D: p << 0.001, n = 80, t_79_ = 17.3). See Figure S2 for transitions between other conditions, and for a comparison of context-specificity with respect to saccade direction.

Since performance in CSG depends on an accurate memory of *t*_*t*_, we hypothesized that these fluctuations may be due to slow drifts in memory. To test this hypothesis, we reasoned that the fluctuations should be smaller if the demands on memory were reduced. Therefore, we trained two monkeys, not previously exposed to the CSG task, to perform a control task in which *t*_*t*_ was measured on every trial, thereby minimizing the need to rely on memory (see Methods). As expected, *t*_*p*_ serial correlations diminished significantly in the control task (Figure S3; p << 0.001, paired sample *t*-test, t_19_ = 4.9, on the partial correlation between CSG and controls, trial lag was limited to 20 trials), suggesting that the slow fluctuations of *t*_*p*_ in the CSG task were a signature of drift in memory. For the remainder of our analyses, we will refer to these fluctuations as memory drifts. Note that this drift reflects a random process with no explicit baseline, and is consistent with a form of memory decay that is characterized by an increasingly broadening of the *t*_*p*_ distribution.

### Reward regulates variability in a context-specific manner

Memory drifts, if left unchecked, could lead to large excursions away from *t*_*t*_, and ruin performance. To maintain a reasonably accurate estimate of *t*_*t*_, another process must counter the drift in memory. In CSG, the only cue that animals can rely on is the trial outcome. Rewarded trials may be used to reinforce the memory while the absence of reward may be construed as evidence that the memory is inaccurate and has to be adjusted. Therefore, we hypothesized that modulations of behavioral variability by reward (Figure 1D) is a signature of an explore-exploit strategy aimed at countering memory drifts.

To test this hypothesis more directly, we defined relative error by *e =* (*t*_*p*_-*t*_*t*_)/*t*_*t*_, and analyzed error statistics on a given trial (*e*^*n*^) as a function of error in the preceding trial (*e*^*n-1*^). We reasoned that errors should have two properties. First, because of the slow memory drifts, errors across consecutive trials should covary; i.e., *μ*(*e*^*n*^) should increase monotonically with *e*^*n-1*^ (Figure 3A, top). Second, an explore-exploit strategy predicts that variability should increase after low or absent reward (Figure 3A, bottom); i.e., *σ*(*e*^*n*^) as a function of *e*^*n-1*^ should have a U-shaped profile (Figure 3A, middle). Indeed, animals’ behavior exhibited both properties: *μ*(*e*^*n*^) increased monotonically with *e*^*n-1*^, and *σ*(*e*^*n*^) had a U-shaped profile (Figure 3B, Figure S4A). We further confirmed these observations statistically using linear regression for *μ*(*e*^*n*^) and quadratic regression for *σ*(*e*^*n*^) (Table 1, see Methods).

**Table 1.**
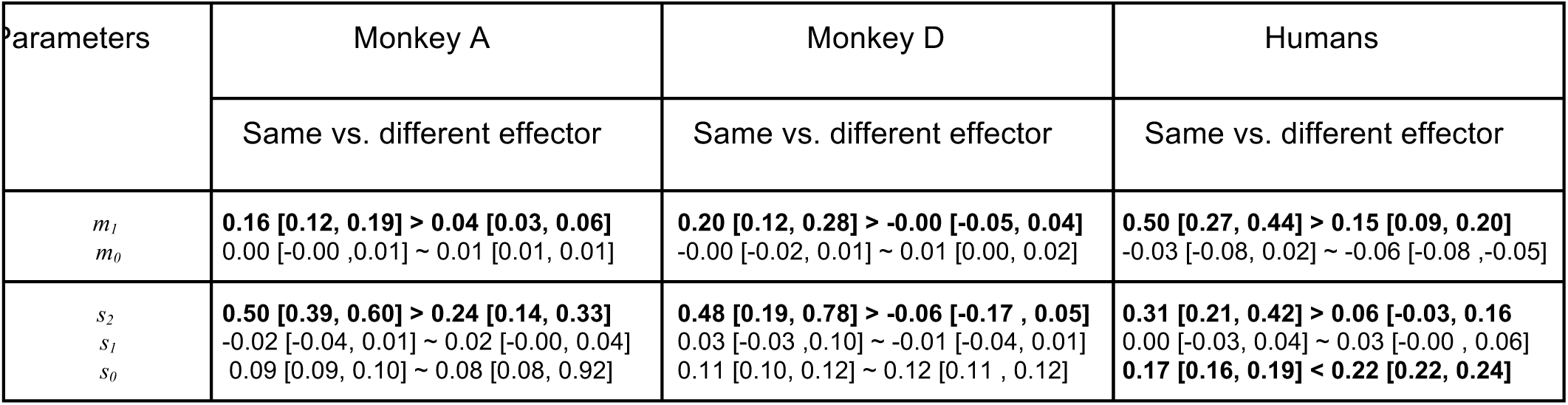
Quantitative assessment of the dependence of *μ*(*e*^*n*^) and *σ*(*e*^*n*^) on *e*^*n-1*^. We used linear regression (*μ*(*e*^*n*^) *= m*_*0*_*+m*_*1*_*e*^*n-1*^) to relate *μ*(*e*^*n*^) to *e*^*n-1*^, and quadratic regression (*σ*(*e*^*n*^) *= s*_*0*_*+s*_*1*_*e*^*n-1*^*+s*_*2*_(*e*^*n-1*^)^*2*^) to relate *σ*(*e*^*n*^) to *e*^*n-1*^. Fit parameters and the corresponding confidence intervals [1%, 99%] are tabulated for each monkey and for humans. We compared the magnitude of fit parameters between the same versus different effector conditions (bold: significantly different).

**Figure 3.**
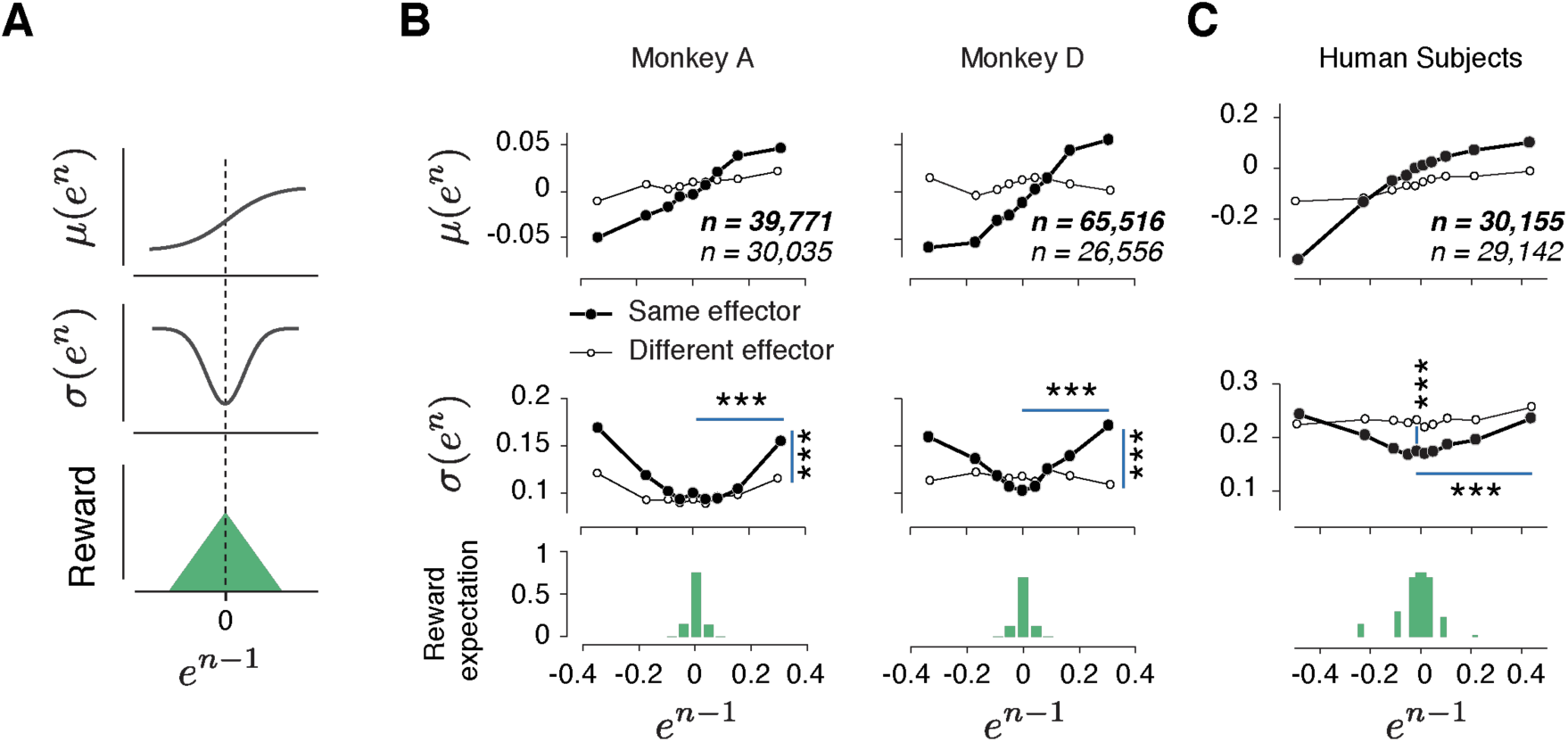
Rapid and context-dependent modulation of behavioral variability by reward. **(A)** Illustration of the expected effect of serial correlations and reward-dependent variability. Top: Positive serial correlations between produced intervals (*t*_*p*_) creates a positive correlation between consecutive errors, and predicts a monotonic relationship between the mean of error in trial *n, μ*(*e*^*n*^), and the value of error in trial *n-1* (*e*^*n-1*^). Middle: Variability decreases with reward. This predicts a U-shaped relationship between the standard deviation of *e*^*n*^, *σ*(*e*^*n*^), and *e*^*n-1*^. Bottom: Reward as a function of *e*^*n-1*^. **(B)** Trial-by-trial changes in the statistics of relative error. Top: *μ*(*e*^*n*^) as a function of *e*^*n-1*^ in the same format shown in (A) top panel. Filled and open circles correspond to consecutive trials of the same and different types, respectively. Middle: *σ*(*e*^*n*^) as a function of *e*^*n-1*^, sorted similarly to the top panel. Bottom: the reward expectation as a function of *e*^*n-1*^. The reward expectation was computed by averaging the magnitude of reward received across trials. In the same effector, variability increased significantly after an unrewarded trials compared to a rewarded trials (horizontal line, two-sample F-test for equal variances, *** p << 0.001) for both large positive errors (Monkey A: F(11169,10512) = 1.09, Monkey D: F(18540,13478) = 1.76) and large negative errors (Monkey A: F(8771,9944) = 1.40, Monkey D: F(21773,14889) = 1.62). The variability after an unrewarded trial of the same effector was significantly larger than after an unrewarded trial of the other effector (vertical line, two-sample F-test for equal variances, *** p << 0.001) for both large positive errors (Monkey A: F(11169,8670) = 1.20, Monkey D: F(18540,7969) = 1.32) and large negative errors (Monkey A: F(8771,5994) = 1.26, Monkey D: F(21773,7179) = 1.27). **(C)** Same as (B) for human subjects. In humans, the variability was also significantly larger after a negatively reinforced trial compared to a positively reinforced trial (horizontal line, two-sample F-test for equal variances, *** p << 0.001) for both large positive errors (F(5536,5805) = 1.19) and large negative errors (F(9366,9444) = 1.11). The variability after a positively reinforced trial of the same effector was significantly lower than after a positively reinforced trial of the other effector (vertical line, two-sample F-test for equal variances, *** p << 0.001, F(14497,15250) = 1.10). For humans, the reward expectation was defined as the ratio between the number of trials with positive feedback and total number of trials.

We performed a number of additional analyses to further scrutinize this finding. For example, we asked whether the modulation of *σ*(*e*^*n*^) could be due to differences across (as opposed to within) behavioral sessions. Applying the same analysis to different grouping of trials verified that the relationship between variability and reward was present within individual behavioral sessions (Figure S4B). Similarly, we asked whether the effect of reward on variability was binary (present/absent) or graded (larger variability for lower reward). More refined binning of trials based on the magnitude of reward provided evidence that the effect was graded: *σ*(*e*^*n*^) was significantly larger in the absence of reward in comparison to low reward, and significantly larger in the low compared to high reward (Figure S4C). These results provide further evidence in support of the relationship between reward and behavioral variability.

Our earlier analysis indicated that the slow memory drifts in the behavior were context-specific (Figure 2B). For the reward modulation of variability to be effective, it must be also context-specific. That is, an increase in variability following a large error should only be present if the subsequent trial is of the same type. Otherwise, reinforcement of one trial type, say ES, may incorrectly adjust variability in the following trials of another type, say HL, and would interfere with the logic of harnessing reward to calibrate the memory of *t*_*t*_. To test this possibility, we analyzed error statistics between pairs of consecutive trials associated with different types (Figure 3B open circles, Figure S4A; Table 1). Results indicated that (1) correlations were strongest between pairs of trials of the same type (as expected from our previous partial correlation analysis in Figure 2A), and (2) the modulation of variability was most strongly and consistently present across trials of the same type. Together, these results provide strong evidence that animals used reward to regulate behavioral variability in the next trial in a context-dependent manner and in accordance with an explore-exploit strategy.

### Reward-dependent context-specific regulation of variability in humans

To ensure that our conclusions were not limited to data collected from highly trained monkeys, we performed a similar experiment in human subjects. In human psychophysical experiment, *t*_*t*_ varied from session to session, and subjects had to constantly adjust their *t*_*p*_ by trial-and-error. Similar to monkeys, human behavior exhibited long-term serial correlations (Figure S6A), and these correlations were context (effector) specific (Figure S6B). We performed the same analysis as in monkeys to characterize the dependence of *μ*(*e*^*n*^) and *σ*(*e*^*n*^) on *e*^*n-1*^. Results were qualitatively similar: *μ*(*e*^*n*^) increased monotonically with *e*^*n-1*^ verifying the presence of slow drifts, and *σ*(*e*^*n*^) had a U-shaped with respect to *e*^*n-1*^ indicating that subjects used the feedback to regulate their variability (Figure 3C and Table 1). Finally, similar to monkeys, the effect of reward on variability was context-specific (Figure 3C). This result suggests that the memory of a time interval is subject to slow drifts and humans and monkeys use reward-dependent regulation of variability as a general strategy to counter these drifts and improve performance.

### Trial outcome causally impacts behavioral variability

So far, our results establish a relationship between trial outcome and behavioral variability. However, since in our experiment error size and trial outcome were deterministically related (larger error led to reduced reward), it is not possible to determine which of these two factors had a causal impact on behavioral variability. More specifically, our current findings are consistent with two models. In one model, trial outcome regulates behavioral variability (Figure 4A; “Causal”). In another, variability is influenced by various upstream factors such as error magnitude, but not by trial outcome (Figure 4A; “Correlational”). To distinguish between these two models, we carried out a new human psychophysics experiment (Figure 4B) in which trial outcome was probabilistic (Figure 4C) so that the presence or absence of a “correct” feedback was not fully determined by the magnitude of the error. This design enabled us to analyze the effect of trial outcome (“correct” versus “incorrect”) on behavioral variability separately for small and large errors.

**Figure 4.**
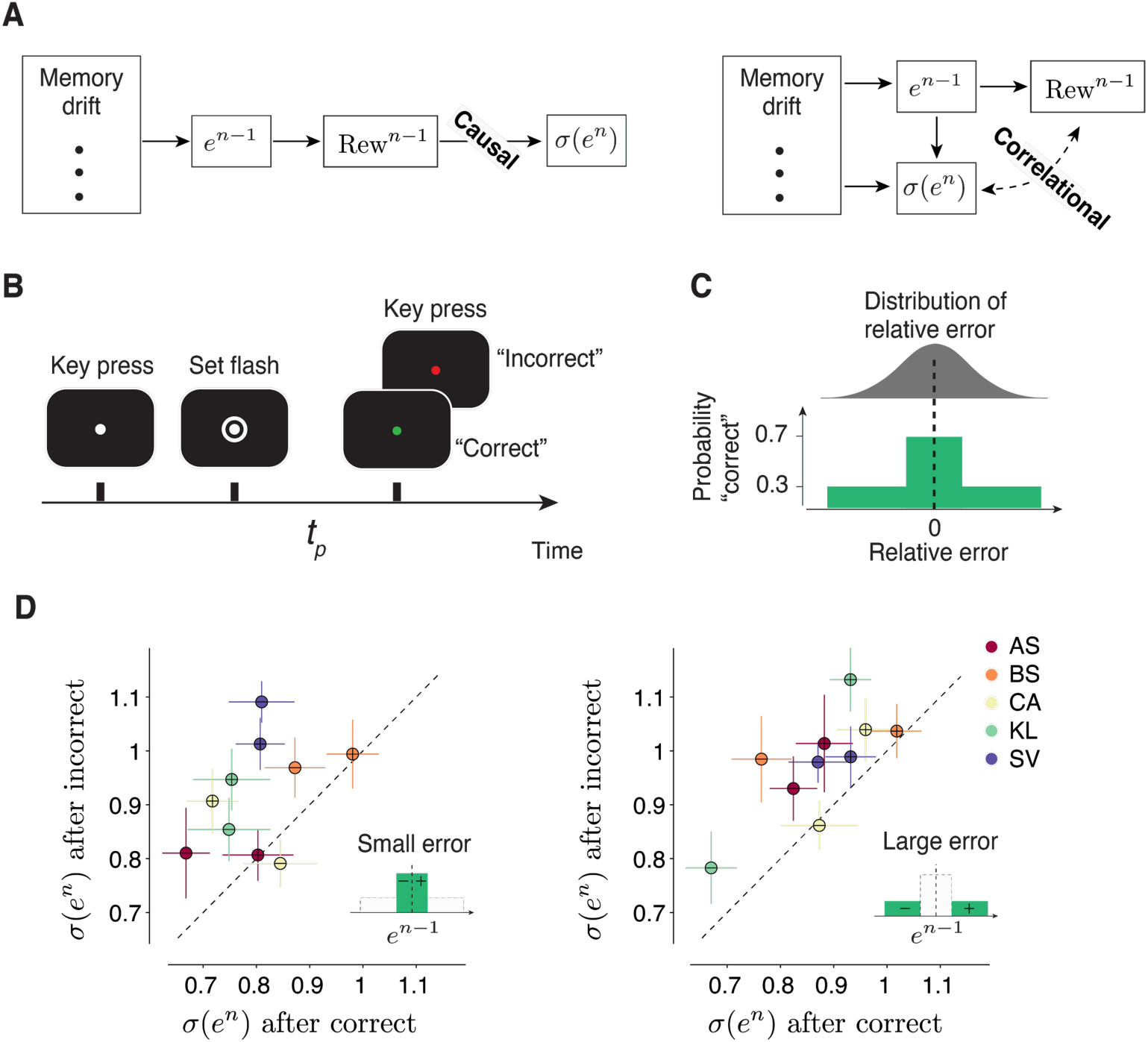
Causal effect of reward on behavioral variability. **(A)** Alternative models of the relationship between the outcome of the preceding trial (Rew^n-1^) and behavioral variability in the current trial (*σ*^*n*^). Left: A model in which Rew^n-1^ causally controls *σ*^*n*^. In this model, various factors (e.g., memory drift) may determine the size of error (*e*^*n-1*^), *e*^*n-1*^ determines Rew^n-1^, and Rew^n-1^ directly regulates *σ*^*n*^. Right: A model in which the relationship between Rew^n-1^ and *σ*^*n*^ is correlational. In this model, Rew^n-1^ is determined by *e*^*n-1*^, and *σ*^*n*^ may be controlled by various factors (including *e*^*n-1*^) but not directly by Rew^n-1^. **(B**,**C)** A time interval production task with probabilistic feedback to distinguish between the two models in (A). **(B)** Trial structure. The subject has to press the spacebar to initiate the trial. During the trial, the subject is asked to hold their gaze on a white fixation spot presented at the center of the screen. After a random delay, a visual ring (‘Set’) is flashed briefly around the fixation spot. The subject has to produce a time interval after Set using a delayed key press. After the keypress, the color of the fixation spot changes to red or green to provide the subject with feedback (green: “correct”, red: “incorrect”). **(C)** Top: A schematic illustration of the distribution of relative error between the produced interval (*t*_*p*_) and the target interval (*t*_*t*_), computed as (*t*_*p*_–*t*_*t*_)/*t*_*t*_. Bottom: After each trial, the feedback is determined probabilistically: the subject is given a “correct” feedback with the probability of 0.7 when *t*_*p*_ is within a window around *t*_*t*_, and with the probability of 0.3 when errors are outside this window. The window length was adjusted based on the overall behavioral variability so that each subject receives approximately 50% “correct” feedback in each behavioral session. **(D)** The causal effect of the outcome of the preceding trial on behavioral variability in the current trial. Left: Scatter plot shows the behavioral variability after “incorrect” (ordinate) versus “correct” (abscissa) trials, for all five subjects, after trials in which *e*^*n-1*^ was small (inset). Results for the positive and negative errors are shown separately (with ‘+’ and ‘-’ symbols, respectively). Right: Same as the left panel for trials in which e^n-1^ was large (inset). In both panels, the variability across subjects was significantly larger following incorrect compared to correct trials (p <<0.001, paired sample t-test, t_199_ = 12.8 for small error, and t_199_ = 13.7 for large error, see Methods).

We compared behavioral variability after “correct” and “incorrect” trials separately for small (Figure 4D, left) and large (Figure 4D, right) errors. Across subjects and irrespective of the size of error, variability was significantly larger after incorrect compared to correct trials (Figure 4D). We also confirmed that the association between variability and trial outcome was not mediated by the size of error (Figure S5C). This result substantiates our hypothesis and rejects all models in which the variability is not explicitly dependent on the trial outcome.

### A generative model linking multiple timescales of variability to reward-based learning

Our results so far indicate that errors in *t*_*p*_ are governed by two key factors: long-term serial correlations creating local and persistent biases in *t*_*p*_, and short-term modulations of *t*_*p*_ variability by reward (see Figure S7 for an analysis of the time lag of the effect of reward). Intuitively, this could provide the means for an efficient control strategy: when bias due to memory drift increases, error increases, reward drops, and the animal seeks to find the correct target interval by increasing variability. Indeed, several recent behavioral experiments have found evidence that is qualitatively consistent with this control strategy (Cashaback et al., 2019; Chen et al., 2017; Dam et al., 2013; Izawa and Shadmehr, 2011; Nikooyan and Ahmed, 2015; Pekny et al., 2015; Shmuelof et al., 2012a; Wu et al., 2014). To assess this idea rigorously, we aimed to develop a generative model that could emulate this control strategy. Initially, we considered two model classes: autoregressive models that can readily capture serial correlations (Wagenmakers et al., 2004) and RL models that use reward to guide behavior (Kaelbling et al., 1996; Sutton and Barto, 1998). However, these two are generally incompatible: classic autoregressive models are insensitive to reward, and RL models cannot accommodate serial correlations that are unrelated to reward. Moreover, RL models are usually geared toward problems with discrete action spaces such as multi-armed bandit problems (Dayan and Daw, 2008), and adaptation of RL to continuous variables using Markovian sampling strategies (Haith and Krakauer, 2014) cannot capture the observed behavioral characteristics (Figure S5A and B).

Based on these results, we concluded that the generative process must have two distinct components, one associated with long-term serial correlations due to the memory drift, and another, with the short-term effect of reward on variability. Accordingly, we sought to develop a Gaussian process (GP) model that could capture both effects. This choice was motivated by three factors: 1) GPs automatically describe observations up to their second order statistics, which are the relevant statistics in our data, 2) GPs offer a nonparametric Bayesian fit to long-term serial correlations, and 3) as we will describe, GPs can be readily augmented to implement short-term control of variability based on reward.

GPs are characterized by a covariance matrix – also known as the GP kernel – that specifies the degree of dependence between samples, and thus determines how slowly the samples fluctuate. The most common and mathematically convenient formulation of this kernel is known as the “squared exponential” kernel function, denoted ***K***_*SE*_ (Figure 5A, top left):

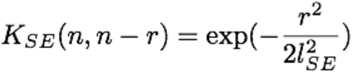

**Figure 5.**
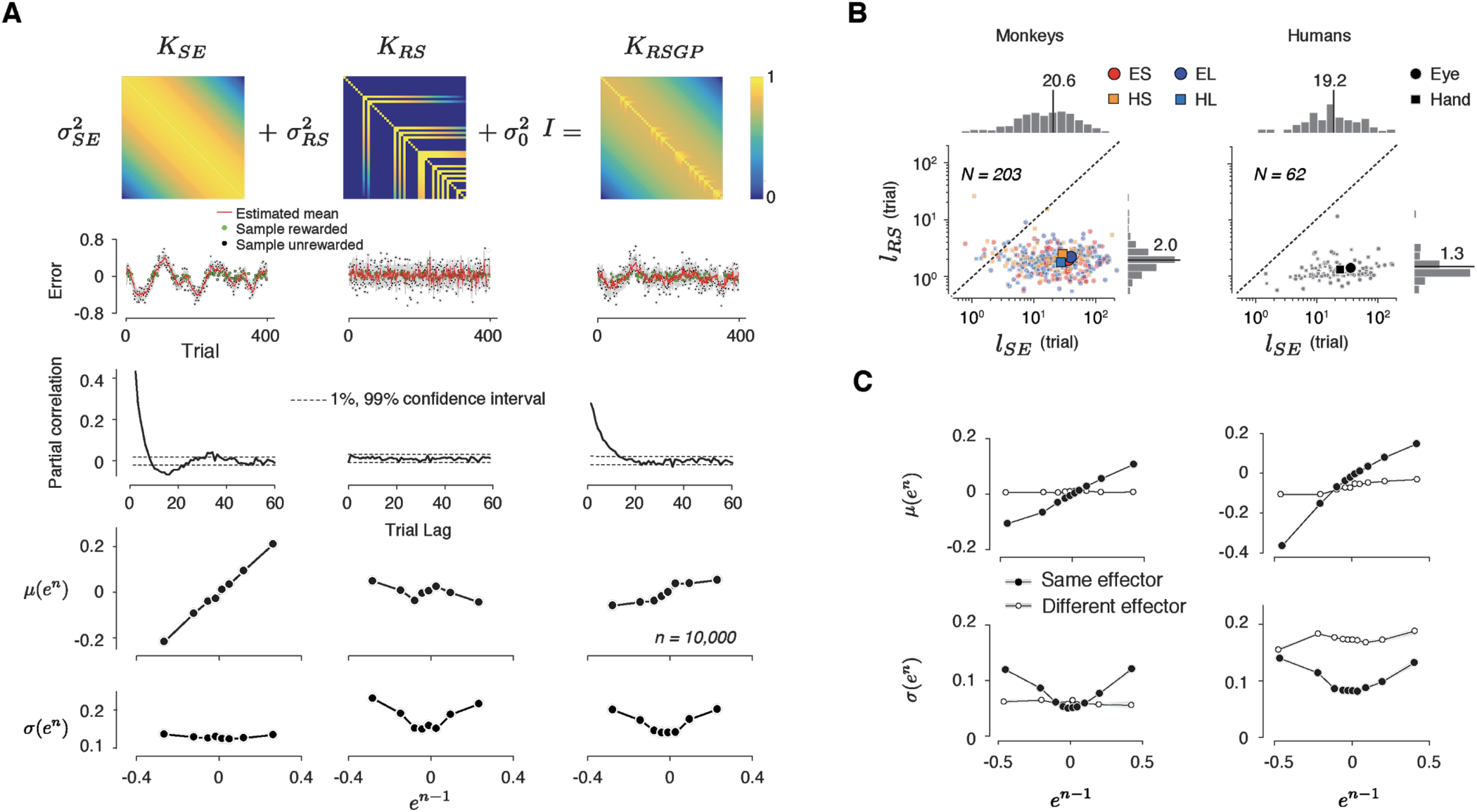
A reward-sensitive Gaussian process model (RSGP) capturing reward-dependent control of variability. **(A)** Top: The covariance function for the RSGP model (***K***_*RSGP*_) is the sum of a squared exponential kernel (***K***_*SE*_), a reward-dependent squared exponential kernel (***K***_*RS*_) and an identity matrix (***I***) weighted by *σ*_*ES*_^*2*^, *σ*_*RS*_^*2*^, and *σ*_*0*_^*2*^, respectively. Second row: Simulations of three GP models, one using ***K***_*SE*_ only (left), one using ***K***_*RS*_ only (middle), and one with the full ***K***_*RSGP*_ (right). Third row: Partial correlation of samples from the three GPs in the second row. Fourth and fifth row: The relationship between the mean (fourth row) and standard deviation (fifth row) of *e*^*n*^ as a function of *e*^*n-1*^ in the previous trial, shown in the same format as in Figure 3B. Only the model with full covariance function captures the observed behavioral characteristics. **(B)** Length scales, *l*_*SE*_ and *l*_*RS*_ associated with ***K***_*ES*_ and ***K***_*RS*_, respectively, derived from fits of RSGP to behavioral data from monkeys (left) and humans (right). Small and large symbols correspond to individual sessions and the median across sessions, respectively. Different trial types are shown with different colors (same color convention as in Figure 1B). *l*_*RS*_ was significantly smaller than the *l*_*SE*_ (monkeys: *p* << 0.001, one-way ANOVA, F_1, 945_ = 463.4; humans: *p* << 0.001, one-way ANOVA, F_1, 235_ = 102.5). **(C)** Statistics of the predicted behavior from the RSGP model fits, shown in the same format as Figure 3B, C. Data were generated from forward prediction of the RSGP model fitted to behavior (see Methods for details). The standard error of the mean computed from n = 100 repeats of sampling of trials is shown as shaded area but it is small and difficult to visually discern.

In ***K***_*SE*_, the covariance between any two samples (indexed by *n* and *n-r*) drops exponentially as a function of temporal distance (*r*) and the rate of drop is specified by the *characteristic length parameter, l*_*SE*_. When *l*_*SE*_ is small, samples are relatively independent, and when it is large, samples exhibit long range serial correlations (Figure 5A, left, 2nd and 3rd row).

Using GPs as the backbone of our model, we developed a reward-sensitive GP (RSGP) whose kernel (***K***_*RSGP*_) is the weighted sum of two kernels, a classic squared exponential kernel (***K***_*SE*_) scaled by *σ*^*2*^_*SE*_, and a reward-sensitive kernel (***K***_*RS*_) scaled by *σ*^*2*^_*RS*_ (Figure 5A, top middle). A third diagonal matrix (*σ*^*2*^_*0*_*I*) was also added to adjust for baseline variance:

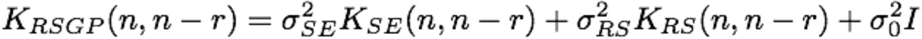

*K*_*RS*_ also has the form of squared exponential with a length parameter, *l*_*RS*_. However, the covariance terms were non-zero only for rewarded trials (Figure 5A, top middle). The reward was a binary variable determined by an acceptance window around *t*_*t*_. This allows rewarded samples to have higher leverage on future samples (i.e., higher covariance), and this effect drops exponentially for trials farther in the future.

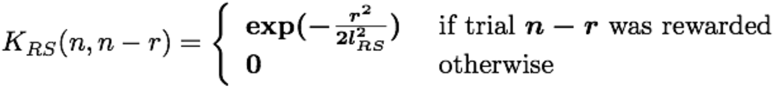

Intuitively, RSGP operates as follows: ***K***_*SE*_ captures the long-term covariance across samples (Figure 5A, 3rd row, left). ***K***_*RS*_ regulates shorter-term covariances (Figure 5A, 3rd row, middle) and allows samples to covary more strongly with recent rewarded trials (Figure 5A, bottom, middle). Non-zero values in ***K***_*RS*_ after rewarded trials strengthen the correlation between samples and lead to a reduction of variability, whereas zero terms after unrewarded trials reduce correlations and increase variability. Using simulations of the full model with ***K***_*RSGP*_ as well as reduced models with only ***K***_*SE*_ or ***K***_*RS*_ (Figure 5A, 2nd row), we verified that both kernels were necessary and that the full RSGP model was capable of capturing the two key features (Figure 5A, 4th and 5th row). Moreover, we used simulations to verify that parameters of the model were identifiable; i.e., fits of the model parameters to simulated data robustly recovered the ground truth (Table S1).

We fitted the RSGP model to behavior of both monkeys and humans (Figure 5B, Figure S8). The recovered characteristic length associated with serial correlations (*l*_*SE*_) were invariably larger than that of the reward-dependent kernel (*l*_*RS*_) (Monkeys: *l*_*SE*_ = 20.6 ± 21.4, *l*_*RS*_ = 2.0 ± 0.7; Humans: *l*_*SE*_ = 19.2 ± 21.8, *l*_*RS*_ = 1.3 ± 0.4; Median ± MAD). The model fit of variances (*σ*_SE_, *σ*_RS_ and *σ*_0_) are shown in Figure S8C. In monkeys, *σ*_0_ and *σ*_SE_ but not *σ*_RS_ were significantly different between two effectors (*σ*_0_: p << 0.001, one-tail two sample t-test, df = 482, t = 5.26; *σ*_SE_: p << 0.001, two sample t-test, df = 482, t = 5.06; *σ*_RS_: p = 0.13, t-test, df = 261, t = 1.5 for the Short interval, and p = 0.26, t-test, dt = 219, t = 1.1 for the Long interval). The dependence of *σ*_0_ and *σ*_SE_ on effector was consistent with our session-wide analysis of variance (Figure 1C). In the human subjects, variance terms were not significantly different between effectors (p = 0.03 for *σ*_0_, p = 0.01 for *σ*_SE_, and p = 0.027 for *σ*_RS_, two sample t-test, df = 118). Importantly, across both monkeys and humans, the model was able to accurately capture the relationship of *μ*(*e*^*n*^) and *σ*(*e*^*n*^) to *e*^*n-1*^ (Figure 5C, and Methods). These results validate the RSGP as a candidate model for simultaneously capturing the slow fluctuations of *t*_*p*_ and the effect of reward on *t*_*p*_ variability.

Two aspects of the RSGP model are noteworthy. First, ***K***_*RS*_ was formulated to capture the effect of reward between trials of the same type, and not transitions between trials associated with different types. Consequently, σ_RS_ for each trial type reflects the effect of reward on variability within the same trial type. The addition of kernels for transitions between effectors was unnecessary because the effect of reward was effector-specific (Figure 3B, Figure S4A). Second, the RSGP was designed to account for the effects of drift and reinforcement regardless of the target interval and thus did not distinguish between the Long and Short conditions. However, the RSGP can be easily reformulated to additionally account for the scaling of variability observed between the Long and Short conditions (Figure S9, and Methods). The RSGP was also able to capture the modulation of behavioral variability in the probabilistic reward experiment (Fig. S5 C and D)

### Slow fluctuations in the cortico-basal ganglia circuits

Recently, we identified a cortico-basal ganglia circuit that plays a causal role in animals’ performance during a flexible timing behavior (Wang et al., 2018). A combination of population data analysis and modeling revealed that animals control the movement initiation time by adjusting a tonic signal in the thalamus that sets the speed at which activity in the dorsomedial frontal cortex (DMFC) and caudate evolve toward an action-triggering state. Based on these results, we hypothesized that memory drifts in behavior may be accompanied by drifts of neural activity in these areas.

To test this hypothesis, we recorded separately from populations of neurons in candidate regions of DMFC, caudate and thalamus. Based on previous work suggesting the importance of initial conditions in movement initiation time (Carpenter and Williams, 1995; Churchland et al., 2006; Hauser et al., 2018; Jazayeri and Shadlen, 2015; Lara et al., 2018; Remington et al., 2018a), we asked whether neural activity near the time of Set (i.e., onset of motor timing epoch) could be used to predict the slow fluctuations in error, which we denote by *e*_slow_ (Figure 6A, Right, red line). To estimate *e*_slow_ on a trial-by-trial basis, we relied on the RSGP model, which readily decomposed the *t*_*p*_ time series into a slow memory-dependent and a fast reward-dependent component. We fitted the RSGP in a context-specific manner and then inferred the value of *e*_slow_ for each trial in a session (see Methods).

**Figure 6.**
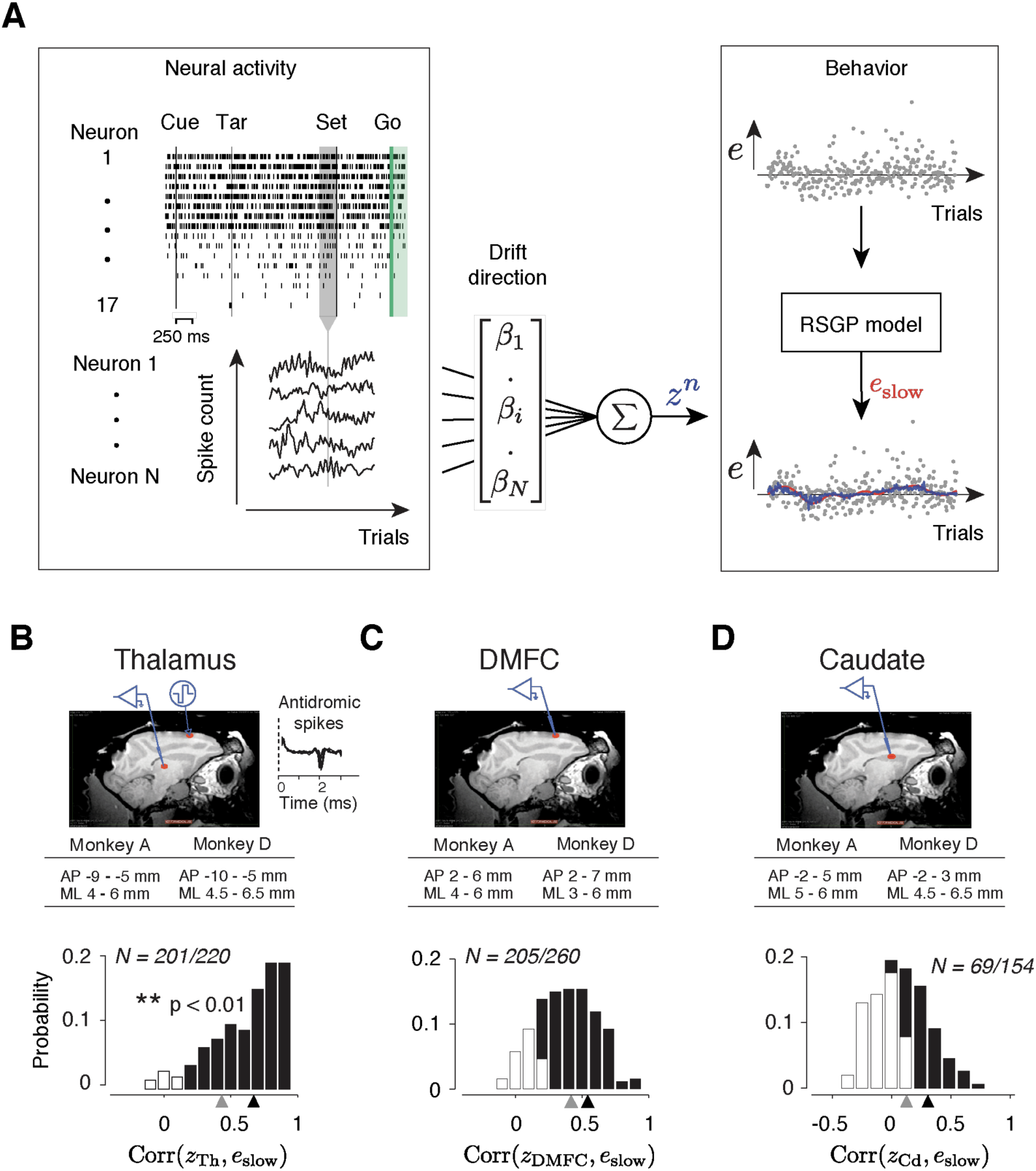
Representation of slow fluctuations of behavior in population activity. **(A)** The schematics of the analysis for identifying the drift direction across a population of simultaneously recorded neurons. Top left: The rows show spike times (ticks) of 17 simultaneously recorded thalamic neurons in an example trial. From each trial, we measured spike counts within a 250 ms window before Set (gray region). Bottom left: The vector of spike counts from each trial (gray vertical line) was combined providing a matrix containing the spike counts of all neurons across all trials. Middle: The spike counts across trials and neurons were used as the regressor in a multi-dimensional linear regression model with weight vector, ***β***, to predict the slow component of error (*e*_*slow*_). Right: We fitted the RSGP to errors (black dots, *e*) and then inferred *e*_*slow*_. The plot shows the neural prediction (*z*^*n*^, blue) overlaid on *e*_*slow*_ derived from RSGP fit to behavior (red line). **(B)** Top: Parasagittal view of one of the animals (monkey D) with a red ellipse showing the regions targeted for electrophysiology. The corresponding stereotactic coordinates relative to the anterior commissure in each animal (AP: anterior posterior; ML: mediolateral). Recorded thalamic neurons were from a region of the thalamus with monosynaptic connections to DMFC(Inset: antidromically activated spikes in the thalamus.) Bottom: Histogram of the correlation coefficients between *e*_slow_ inferred from the RSGP model and *z*^*n*^_Th_ (projection of thalamic population activity on drift direction) across recording sessions. Note that some correlations are negative because of cross-validation Black bars correspond to the sessions in which the correlation was significantly positive (**p < 0.01; hypothesis test by shuffling trial orders). The average correlation across all sessions and the average of those with significantly positive correlations are shown by gray and black triangles, respectively. **(C)** Same as B for DMFC. **(D)** Same as B for the caudate.

We measured spike counts within a 250-ms window before Set (Figure 6A, Left), denoted ***r***, and formulated a multi-dimensional linear regression model to examine the trial-by-trial relationship between ***r*** and *e*_slow_. We computed the regression weight, ***β***, that when multiplied by ***r***, would provide the best linear fit to *e*_slow_. Considering the vector of spike counts in each trial as a point in a coordinate system where each axis corresponds to the activity of one neuron (“state space”), ***β*** can be viewed as the direction along which modulations of spike counts most strongly correlate with memory drifts. Accordingly, we will refer to the direction associated with ***β*** as the drift direction, and will denote the strength of activity along that direction by *z* (*z* = ***rβ***, Figure 6A, Right, blue line).

To test the hypothesis that memory drifts were accompanied by drifts of neural activity in the thalamus, DMFC and caudate, we used a cross-validated procedure (see Methods) to compute ***β***, derive a trial-by-trial estimate of *z*, and measure the correlation between *z* and *e*_slow_ on a session-by-session basis. We found strong evidence that neural activity in all three areas exhibited slow fluctuations in register with memory drifts. In 91%, ~79%, and ~45% of sessions, the activity along the drift direction in the thalamus (*z*_Th_), DMFC (*z*_DMFC_), and caudate (*z*_Cd_), respectively, was significantly correlated with *e*_slow_ (Figure 6B-D, p < 0.01, null hypothesis test by shuffling trials).

### Reward-dependent regulation of variability in the thalamus

Next, we analyzed the statistics of neurally-inferred drift (*z*) across pairs of consecutive trials using the same approach we applied to error statistics in behavioral data (Figure 3B). To do so, we extracted pairs of (*e*^*n-1*^, *z*^*n*^) for consecutive trials of the same type and binned them depending on the value of *e*^*n-1*^. Since all three areas carried a signal reflecting memory drifts, we expected *μ*(*z*^*n*^) in all the areas to have a monotonic relationship with *e*^*n-1*^. Results were consistent with this prediction as evidenced by the slope of a linear regression model relating *μ*(*z*^*n*^) to *e*^*n-1*^ (filled circles in Figure 7B-D top, Table 2). Moreover, this relationship was absent for consecutive trials associated with different effectors (open circles in Figure 7B-D top, Table 2). These results substantiate the presence of context-specific fluctuations of neural activity in register with drifts in animals’ memory of *t*_*t*_ across populations of neurons in the thalamus, DMFC and caudate.

**Table 2.**
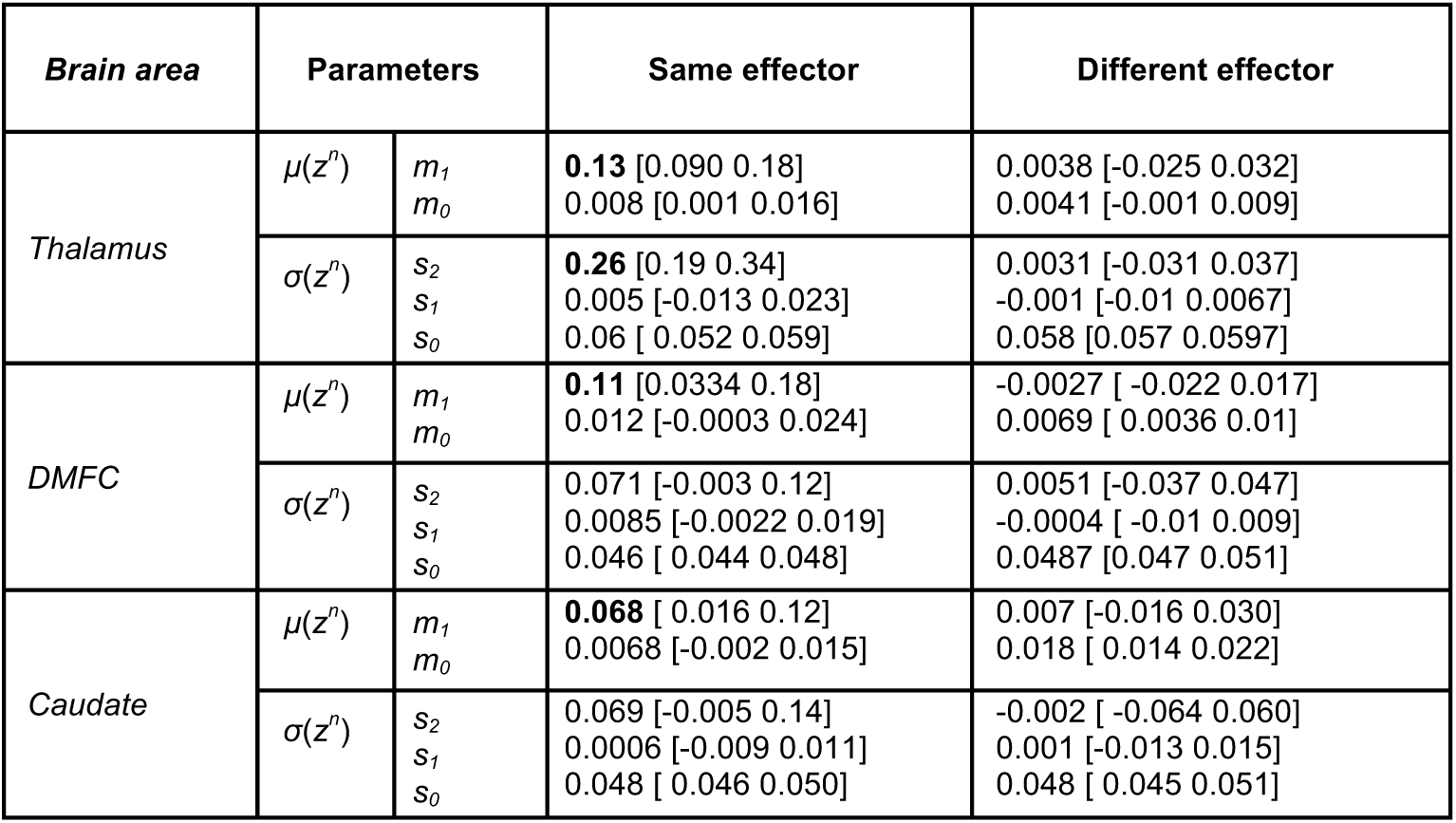
Regression model fits relating spike count along the drift direction on trial *n* (*z*^*n*^) to error in trial *n-1* (*e*^*n-1*^). *m*_*0*_ and *m*_*1*_ are parameters of the linear regression model relating the mean of *z*^*n*^ (*μ*(*z*^*n*^)) to *e*^*n-1*^; i.e., *μ*(*z*^*n*^) *= m*_*0*_*+m*_*1*_*e*^*n-1*^. *s*_*0*_, *s*_*1*_ and *s*_*2*_ are parameters of the quadratic regression model relating the standard deviation of *z*^*n*^ (*σ*(*z*^*n*^)) to *e*^*n-1*^; i.e., *σ*(*z*^*n*^) *= s*_*0*_*+s*_*1*_*e*^*n-1*^*+s*_*2*_(*e*^*n-1*^)^*2*^. Fit parameters are shown separately for the thalamus, DMFC and caudate and further separated depending on whether trial *n-1* and *n* were of the same or different effectors. Bold values for *m*_*1*_ and *s*_*2*_ were significantly positive (** p < 0.01, 1% and 99% confidence intervals of the estimation were shown).

**Figure 7.**
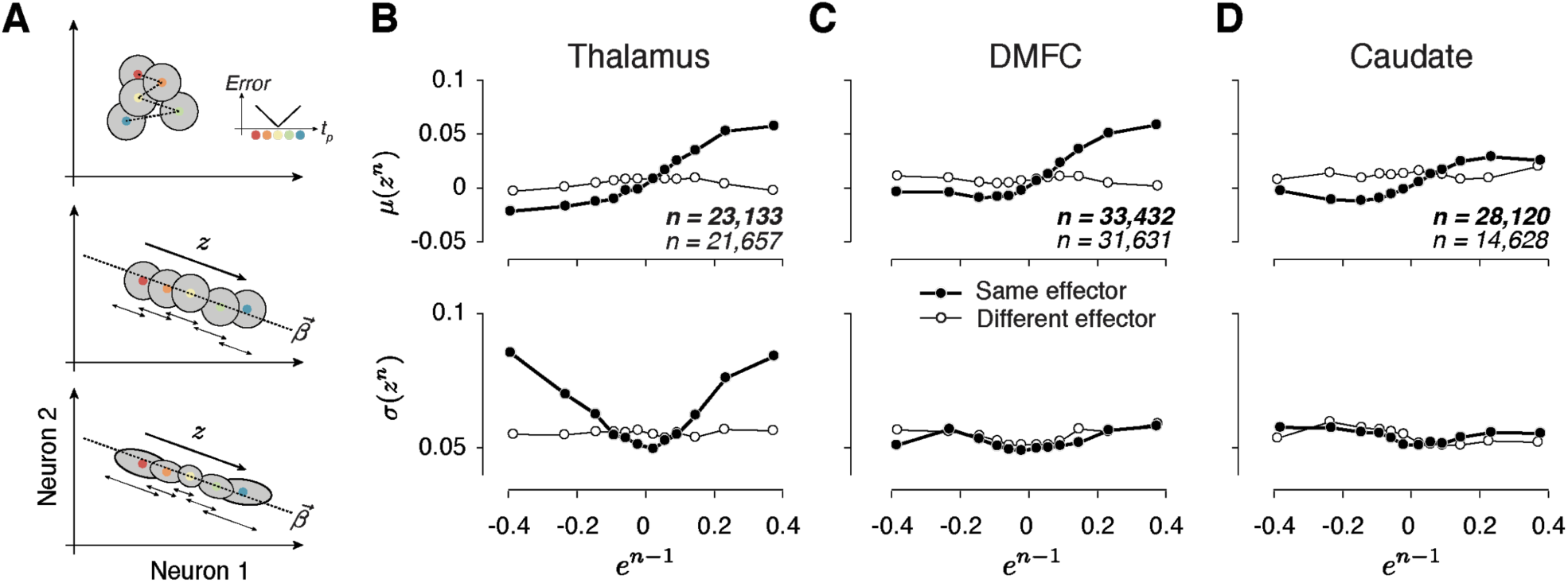
Alignment of reward-dependent neural variability and drift in thalamus, but not in DMFC and caudate. **(A)** Various hypotheses for how population neural activity could be related to produced intervals (*t*_*p*_) shown schematically in 2 dimensions (2 neurons). Top: The average neural activity (colored circles) is not systematically organized with respect to *t*_*p*_, and the trial-by-trial variability of spike counts for a given *t*_*p*_ around the mean (gray area) is not modulated by the size of the error. The inset shows how error increases as *t*_*p*_ moves away from the target interval (*t*_*t*_). Middle: Projection of average activity along a specific dimension (dotted line) is systematically ordered with respect to *t*_*p*_, but the variability (small stacked arrows) does not depend on the size of the error. Bottom: Projection of average activity along a specific dimension is systematically ordered with respect to *t*_*p*_ and the variability along the same axis increases with the size of error. **(B)** The relationship between neural activity in the thalamus on trial *n* and relative error in the preceding trial (*e*^*n-1*^). Top: Expected mean of population activity on trial *n* (*μ*(*z*^*n*^)) along the drift direction (***β***) as a function of *e*^*n-1*^. Bottom: Expected standard deviation of population activity along the drift direction on trial *n* (*σ*(*z*^*n*^)) as a function of *e*^*n-1*^. Results are shown in the same format as in Figure 3B (thick lines: same effector; thin lines: different effectors). **(C)** and **(D)** Same as (B) for population activity in DMFC and caudate. See Figure S10 for the result of individual animal.

A critical question was whether, in any of these areas, reward regulates the variability of neural activity. Importantly, the reward-dependent modulation of variability should be restricted to the drift direction in the population activity (Figure 7A, bottom); otherwise, this strategy will not be able to effectively counter the degrading effect of memory drift in a context-dependent manner. In thalamus, *σ*(*z*^*n*^_Th_) exhibited the characteristic U-shape profile with respect to *e*^*n-1*^ (Figure 7B bottom), which we verified quantitatively by comparing the variance of *z*^*n*^_Th_ for rewarded and unrewarded trials (p < 0.01, two-sample F-test for equal variances on *z*^*n*^_Th_, F(5608,4266) = 1.08 for negative *e*^*n-1*^ and p < 0.001 for positive *e*^*n-1*^, F(4723,6125) = 1.28), and by testing the profile using quadratic regression (see Methods; Table 2).

As an important control, we performed the same analysis between consecutive trials associated with different effectors and we found no significant relationship between *σ*(*z*^*n*^_Th_) and *e*^*n-1*^ (p = 0.91, two-sample F-test for equal variances, F(5332,3984) = 1.0 for the negative *e*^*n-1*^; p = 0.97 for the positive *e*^*n-1*^, F(4234,5857) = 0.99). Note that these analyses were repeated after square-root transforming spike count data to stabilize Poisson-like variability; this did not affect any of our conclusions. Taken together, these results indicate that variability of thalamic responses was modulated by reward, and that this modulation was aligned to the drift direction.

Unlike the thalamus, in DMFC and caudate, variability along the memory drift was relatively independent of *e*^*n-1*^ (Figure 7C bottom, 6D bottom); i.e., *σ*(z^*n*^_DMFC_) and *σ*(z^*n*^_Cd_) were not significantly different after rewarded and unrewarded trials (two-sample F-test for equal variances, F(6244,4818) = 0.87, p = 0.99 for the negative *e*^*n-1*^; F(4825,7572) = 1.002, p = 0.021 for the positive *e*^*n-1*^, see Figure S10 for each animal separately). We verified the absence of a U-shape profile using quadratic regression (see Methods; Table 2) and ensured that the lack of modulation in DMFC and caudate compared to the thalamus was not because of a difference in the number of simultaneously recorded neurons (Figure S11 B and C). Together, these results provide evidence that the effect of reward in DMFC and caudate was not aligned to the ***β*** associated with the slow fluctuations. We note, however, that an unconstrained decoding strategy aimed at simultaneously capturing both components of errors can find directions along which the slow fluctuations and the effect of reward are aligned in all three areas (Figure S12).

Our analysis so far focused on spike counts in a fixed 250-ms time window before Set. However, the effect of reward on variability in the thalamus (absent in DMFC and caudate) was present throughout the trial, and extended as far back as the preceding feedback (Figure 8A top; *** p <0.001, two-sample F-test for equal variances). Together, these results suggest that reward exerts its influence on behavior by regulating the variability of thalamic signals driving movement initiation time.

**Figure 8.**
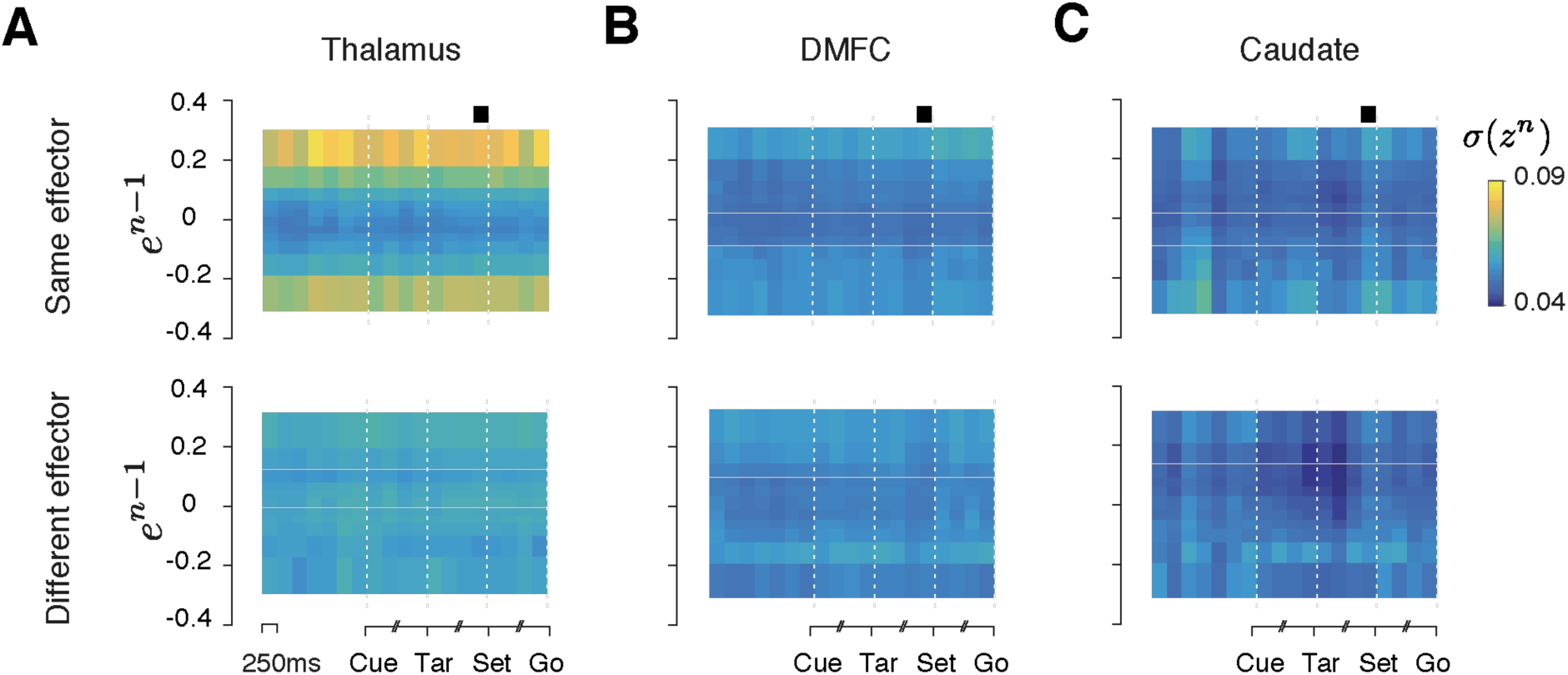
Reward-dependent neural variability throughout the trial. **(A)** Average standard deviation of population activity in the thalamus at different points throughout the trial aligned to the Cue, Tar, Set and Go events. For each time point, we inferred the drift direction using the same analysis shown in Figure 6A, and projected spike counts onto the drift direction. We denote the projection of trial *n* by *z*^*n*^. Each column shows the standard deviation of *z*^*n*^ (*σ*(*z*^*n*^)) as a function of error in the preceding trial (*e*^*n-1*^) based on spike counts within a 250 ms window centered at a particular time point in the trial. Black square: the time window used for the rest of this paper. We grouped *e*^*n-1*^ in to 12 bins, 6 for negative *e*^*n-1*^ and 6 for positive *e*^*n-1*^. Results are shown separately for conditions in which tials *n-1* and *n* were of the same or different effectors. (*** p< 0.001, using a two-sample F-test comparing the variance of *z*^*n*^ between rewarded and unrewarded trials separately for positive and negative values of *e*^*n-1*^; H0: equal variance for at least one of the comparisons). **(B)** and **(C)** Same as (A) for DMFC and caudate. Results are for data combined across two animals. Figure S13 shows the results of the same analysis for each animal separately.

## Discussion

Variability is a ubiquitous property of behavior that can either degrade performance or promote learning through exploration. Here, we were able to advance our understanding of the function and neurobiology of variability in three directions. First, we found an important role for memory drifts in timing variability, and showed that humans and monkeys use reward to calibrate memory against such drifts. Second, using model-based analysis of behavior, we showed that reward promotes learning through adjustment of variability. Finally, we characterized the neural underpinnings of variability associated with memory drifts and reinforcement across populations of neurons within the cortical-basal ganglia circuits.

### Role of memory drift in behavioral variability

A key feature of the behavioral data was the presence of slow fluctuations in produced intervals leading to serial correlations extending over minutes and across dozens of trials. These fluctuations have been reported in various behavioral tasks (Chen et al., 1997; Gilden et al., 1995; Murakami et al., 2017), and are typically attributed to fatigue, arousal or other nonspecific factors modulating internal states (Harris and Thiele, 2011; Kato et al., 2012; Lee and Dan, 2012; Luck et al., 1997; Niell and Stryker, 2010; Vinck et al., 2015). Although slow internal state modulations are likely to present in our experiment, they cannot be the only driver since the serial correlations were strongly context-specific. Based on these observations, we reasoned that these fluctuations may reflect drifts in memory. This interpretation was consistent with the results of our control experiment showing diminished serial correlations when memory demands were reduced (Figure S3). However, further work is needed to fully characterize the nature of these slow fluctuations. For example, the slow component may be in part a reflection of an active averaging process to maintain a stable memory (Joiner and Smith, 2008), as suggested by error-based motor learning studies (Huberdeau et al., 2015; Smith et al., 2006; Wolpert et al., 2011).

A more puzzling observation was the specificity of memory drifts with respect to the target interval for the same effector. We currently do not have a definite explanation for this result, but our previous work in the domain of time interval production (Wang et al., 2018) and reproduction (Remington et al., 2018a) as well as others studies in the motor system (Afshar et al., 2011; Ames et al., 2014; Hauser et al., 2018; Sheahan et al., 2016; Vyas et al., 2018) suggest that several aspects of movement control can be understood in terms of adjusting inputs and initial conditions of a dynamical system (Churchland et al., 2012; Remington et al., 2018b). Accordingly, the interval specificity of the memory drifts suggests that distinct patterns of neural activity set the interval-dependent input and/or initial condition, which is consistent with our previous work (Wang et al., 2018).

### Role of reinforcement in behavioral variability

In our task, reinforcement is the only information provided experimentally that subjects can use to calibrate their memory of the target interval. Therefore, the computational demands in our task fall squarely within the framework of reinforcement learning (RL). Most existing RL models have focused on experimental settings in which the agent faces a discrete set of options and/or a limited action space(Daw et al., 2006; Hayden et al., 2011; Lee et al., 2011; Wilson et al., 2014). In these situations, RL models posit that the agent keeps track of the value of available options and adopts a suitable policy to choose among them (Sutton and Barto, 1998). In our task, subjects have to use reinforcement to choose the correct interval, which is a continuous variable. In theory, RL can be extended to continuous variables, but doing so would require the brain to represent an infinite-dimensional value function, which seems implausible. Moreover, in this case, noise associated with producing an interval would cause the produced interval to differ from what was intended, and that would interfere with the agent’s ability to correctly evaluate the intended action.

Due to these complications, several recent studies have proposed an alternative explore-exploit strategy for continuous variables in which the agent uses reinforcement to directly regulate behavioral variability (Pekny et al. 2015; Santos et al. 2015; Wu et al. 2014; Dhawale et al. 2017; Fee and Goldberg 2011; Tumer and Brainard 2007). Our results were consistent with this view: variability was modulated by reward in a manner that was consistent with an explore-exploit strategy. This hypothesis was strongly supported by the results of our psychophysical experiment using a probabilistic reward schedule, which confirmed that the effect of trial outcome on behavioral variability was causal and independent of the magnitude of error (Figure 4). Another key observation was that the effect of reward on behavioral variability was context-specific (Figure 3); i.e., variability associated with producing a specific interval with a specific effector was most strongly dependent on reward history in trials of the same interval and effector. Given that the memory drifts were also context-specific, this finding indicates that one important function of reward-dependent regulation of variability is to counter memory drifts.

### Extending reinforcement learning to continuous variables

Reinforcement learning through reward-dependent modulation of behavioral variability is an appealing idea that is gaining prominence, especially within the domain of motor learning (Dhawale et al., 2017). Here, we developed a reward-sensitive Gaussian process (RSGP) model to rigorously characterize and quantify this learning process. RSGP was able to capture both the long-term effect of memory drift, and the short-term effect of reward on variability. It also captured behavioral observations in the probabilistic reward experiment where we validated the causal effect of reward on variability (Figure S5C, D). A recent motor learning study in rodents found that the effect of reward on behavioral variability may last a few trials into the future (Dhawale et al. 2019). This was also evident in our data (Figure S7), and RSGP provided a quantitative measure of the temporal extent of this effect (Figure S8). As such, we expect RSGP to help future studies quantify the strength and persistence with which reinforcement guards against ubiquitous nonstationarities in behavior (Gilden et al. 1995; Weiss et al. 1955; Merrill and Bennett 1956; Laming 1979; Chaisanguanthum et al. 2014; Murakami et al. 2017; Chen et al. 1997).

### Memory drift in the cortico-basal ganglia circuits

Behavioral variability in timing tasks is thought to have a multitude of distributed biophysical and synaptic origins (Gibbon et al., 1984; Mauk and Buonomano, 2004; Paton and Buonomano, 2018). Several studies have been able to trace this variability *in-vivo* to spiking activity of neurons in cortico-basal ganglia circuits (Murakami et al. 2017; Murakami et al. 2014; Gouvêa et al. 2015; Merchant and Averbeck 2017; Wang et al. 2018; Dhawale et al. 2017). We previously found that a circuit comprised of DMFC, DMFC-projecting thalamus and caudate plays a causal role in the control of movement initiation time (Wang et al., 2018), and that certain patterns of activity in each area were correlated with behavioral variability on a trial-by-trial basis. Here, we additionally confirmed that population activity in these areas carries a signal correlated with slow fluctuations of behavior. This finding is broadly consistent with previous studies reporting correlates of internal state changes and/or slow behavioral fluctuations in the thalamus (Halassa et al., 2014), the medial frontal cortex (Karlsson et al., 2012; Murakami et al., 2017; Narayanan and Laubach, 2008; Sul et al., 2010), and the caudate (Lau and Glimcher, 2007; Lauwereyns et al., 2002; Santos et al., 2015). However, the effector- and interval-dependent nature of these fluctuations in our data suggests that they may partially reflect context-specific memory drifts. The key feature of our task that enabled us to uncover this specificity was the alternation between different contexts on a trial-by-trial basis. Since many previous studies did not include this feature, it is possible that certain aspects of neural variability previously attributed to nonspecific internal state changes were in part caused by memory drifts related to specific task rules and parameters. Indeed, drifts and degradation of instrumental memories may be a key limiting factor in motor skill performance (Ajemian et al., 2013).

Although we found a correlate of these drifts in all three brain areas, we cannot make a definitive statement about the loci of the underlying synaptic and biophysical drifts. It is likely that the memory has a distributed representation, in which case the drift may result from stochastic processes distributed across multiple brain areas. It is also possible that different contexts engage specific sub-circuits such as corticostriatal synapses (Fee and Goldberg, 2011; Xiong et al., 2015) and circuit-level interactions allow these drift to be manifest across other nodes of the cortico-basal ganglia circuit.

For different effectors, context-specificity may be attributed to different patterns of activity across the same population of neurons or by distinct populations of neurons. Our results in the cortico-basal ganglia circuits provided evidence for the former (Figure 7, S15). However, we cannot rule out the possibility that this context-specificity arises partially from execution noise within distinct effector-specific downstream areas (e.g, brainstem) as we did not record from those areas. However, execution noise is generally considered to be irreducible (Dhawale et al., 2017; Faisal et al., 2008), and is therefore not expected to be influenced by reinforcement. Previous work suggests that central noise (e.g., in the cortico-basal ganglia circuits) plays a significant role in output variability (van Beers, 2009; Churchland et al., 2006). It is this portion of variability that is likely subject to adjustments through reinforcement.

### Reinforcement via thalamus

Next, we asked whether the variability of neural activity in the thalamus, DMFC and caudate was modulated by reward in the preceding trial in the same context-dependent manner as in the behavior. According to our hypothesis, the key function of the reward-dependent regulation of variability is to counter the memory drifts. This hypothesis makes a specific prediction: reward should modulate the specific pattern of population neural activity that corresponds to memory drifts in the behavior, which we referred to as drift direction. Analysis of neural activity revealed that this effect was present in the thalamus but not in DMFC or caudate. In the thalamus, spike count variability along the drift direction increased after rewarded trials and decreased after unrewarded trials in a context-specific manner. Previous studies have reported that in the thalamus firing rates are modulated on a trial-by-trial basis by attention (Mcalonan et al., 2008; Saalmann et al., 2012; Zhou et al., 2016) and rule/context-dependent computations (Schmitt et al., 2017; Wang et al., 2018). Our work demonstrates that modulation of thalamic activity may additionally subserve reward-based calibration of movement initiation times. It will be important for future studies to investigate whether this finding generalizes to other movement parameters.

The fact that the same effect was not present in DMFC and caudate serves as a negative control and thus strengthens our conclusions. However, this begs the question of why this regulation was not inherited by DMFC and caudate, especially given that DMFC receives direct input from the region of the thalamus we recorded from. The answer to this question depends on the nature of signal transformations along the thalamocortical pathway. While some experiments have suggested similar response properties for thalamus and their cortical recipients (Guo et al., 2017; Sommer and Wurtz, 2006), others have found that thalamic signals may undergo specific transformations along the thalamocortical pathway (Berman and Wurtz, 2011; Hubel and Wiesel, 1962; Schmitt et al., 2017; Wang et al., 2018; Wimmer et al., 2015). Therefore, the extent to which we should expect responses in DMFC and thalamus to have similar properties is unclear.

Our approach for investigating whether the effect of reinforcement was aligned with memory drift was correlative: we used a cross-validated decoding strategy to infer the response pattern most strongly associated with memory drift and tested whether reinforcement exerted its effect along that pattern. According to this analysis, only in thalamus the two effects were appropriately aligned. However, we cannot rule out the possibility that behavior was controlled by a pattern of activity different from those we inferred from our analysis. Indeed, with a complementary analysis using an unconstrained decoding approach, we were able to find patterns of activity in all three brain areas that simultaneously reflected the effects of memory drift and reinforcement (Figure S12). Therefore, an important consideration for future work is to use perturbation methods to verify causally the direction in the state space associated with memory drifts and reinforcement.

However, other considerations are consistent with thalamus playing a strong role in controlling timing variability. For example, it has been shown that inactivation of thalamus has a much stronger effect on modulating animals’ motor timing variability compared to DMFC and caudate (Wang et al., 2018). Moreover, it has been shown that the nature of signals in DMFC-projecting thalamus and DMFC during motor timing are indeed different: DMFC neurons have highly heterogeneous response profiles that evolved at different speeds depending on the interval, whereas thalamic neurons carried signals whose strength (i.e., average firing rate) encoded the underlying speed. This transformation may provide an explanation for why reward-dependent modulation of firing rates was evident in the thalamus but not in DMFC. Since thalamic neurons encode interval in their average firing rates, it is expected that regulation of timing variability by reward would similarly impact average firing rates. In contrast, in DMFC, the key signature predicting behavior was the speed at which neural trajectories evolved over time – not the average firing rates. This predicts that reward should alter the variability of the speed of neural trajectories. In principle, it is possible to verify this prediction by estimating the variance of the speed of neural trajectories as a function of reward. However, this estimation is challenging for two reasons. First, speed in a single trial is derived from changes in instantaneous neural states, and the estimation of instantaneous neural states is unreliable unless the number of recorded neurons exceeds the dimensionality of the subspace containing the neural trajectory (Gao et al., 2017). Second, our predictions are about the variance – not mean – of speed, and estimating variance adds another layer of statistical unreliability unless the number of neurons or trials are sufficiently large.

Nonetheless, we devised a simple analysis to estimate the variance of the speed of neural trajectories across single trials in all three areas. Results were consistent with our predictions: variance of speed was larger after unrewarded trials in DMFC and caudate but not thalamus, and this effect was present only for consecutive trials associated with the same effector (Figure S16). In other words, our results agree with the interpretation that reward controls the variance of average firing rates in the thalamus, and this effect leads to the control of the variance of the speed at which neural trajectories evolve in DMFC and caudate.

One open question pertains to which brain areas supply the relevant information for the reward-dependent control of behavioral variability. While we cannot address this question definitively, we note that the area of thalamus we have recorded from receives information from three major sources, the frontal cortex, the output nuclei of the basal ganglia, and the deep nuclei of the cerebellum (Kunimatsu et al., 2018; Middleton and Strick, 2000). Modulation of variability prior to movement initiation has been reported in motor and premotor areas (Churchland et al., 2006, 2010), and distinct correlates of explore-exploit strategy has been found across multiple nodes of the frontal cortex (Ebitz et al., 2018; Hayden et al., 2011; Massi et al., 2018; Sarafyazd and Jazayeri, 2019; Tervo et al., 2014). Therefore, it is possible that modulation of variability in thalamus originates in the cortex. One salient example where cortical variability can be adjusted rapidly and in a behaviorally-relevant fashion is in the domain of attentional control where spatial cueing can lead to a drop of correlated variability across neurons whose receptive fields correspond to the cued location (Cohen and Maunsell, 2009; Mitchell et al., 2009; Ni et al., 2018; Ruff and Cohen, 2014). It is possible that a similar strategy is at play in the frontal cortex in the service of exploratory learning. However, to act as an effective learning mechanism, such correlated cortical variability must be additionally sensitive to reward-dependent neuromodulatory signals such as dopamine (Frank et al., 2009) possibly by acting on local inhibitory neurons (Huang et al., 2019). The basal ganglia could also play a role in reward-dependent control of thalamic firing rates (Kunimatsu and Tanaka, 2016; Kunimatsu et al., 2018). For example, single neuron responses in substantia nigra pars reticulata that are strongly modulated by reward schedule (Yasuda and Hikosaka, 2015) can influence neural responses in the thalamus. Finally, the cerebellum plays a central role in trial-by-trial calibration of motor variables (Herzfeld et al., 2015; Ito, 2002; Medina and Lisberger, 2008) including movement initiation time (Ashmore and Sommer, 2013; Kunimatsu et al., 2018; Narain et al., 2018) and thus is a natural candidate for calibrating firing rates in thalamus although how such calibration could be made reward-sensitive remains an open question (Hoshi et al., 2005). In sum, our work provides behavioral, modeling and neurophysiological evidence in support of the hypothesis that the brain uses reinforcement to regulate behavioral variability in a context-dependent manner.

## Materials and Methods

Two adult monkeys (Macaca mulatta; one female, one male), and five human subjects (18-65 years, two females and three males) participated in the main task. Two monkeys (both male) participated in the memory control experiment. In addition, five more human subjects (18-65 years, two females and three males) participated in the probabilistic reward task to test the causal effect of reward on variability. The Committee of Animal Care and the Committee on the Use of Humans as Experimental Subjects at Massachusetts Institute of Technology approved the animal and human experiments, respectively. All procedures conformed to the guidelines of the National Institutes of Health.

### Animal experiments

Monkeys were seated comfortably in a dark and quiet room. The MWorks software package (https://mworks.github.io) running on a Mac Pro was used to deliver stimuli and to control behavioral contingencies. Visual stimuli were presented on a 23 inch monitor (Acer H236HL, LCD) at a resolution of 1920×1080, and a refresh rate of 60Hz). Auditory stimuli were played from the computer’s internal speaker. Eye position was tracked with an infrared camera (Eyelink 1000; SR Research Ltd, Ontario, Canada) and sampled at 1 kHz. A custom-made manual button, equipped with a trigger and a force sensor, was used to register button presses.

#### The Cue-Set-Go task

Behavioral sessions in the main experiment consisted of four randomly interleaved trial types in which animals had to produce a target interval (*t*_*t*_) of either 800 ms (Short) or 1500 ms (Long) using either a button press (Hand) or a saccade (Eye). The trial structure is described in the main Results (Figure 1A). Here, we only describe the additional details that were not described in the Results. The “Cue” presented at the beginning of each trial consisted of a circle and square. The circle had a radius of 0.2 deg and was presented at the center of the screen. The square had a side of 0.2 deg and was presented 0.5 deg below the circle. For the trial to proceed, the animal had to foveate the circle (i.e., eye fixation) and hold its hand gently on the button (i.e., hand fixation). The animal had to use the hand contralateral to the recorded hemifield. We used an electronic window of 2.5 deg around the circle to evaluate eye fixation, and infrared emitter and detector to evaluate hand fixation. After 500-1500 ms delay period (uniform hazard), a saccade target was flashed eight deg to the left or right of the circle. The saccade target (“Tar”) had a radius of 0.25 deg and was presented for 250 ms. After another 500-1500 ms delay (uniform hazard), an annulus (“Set”) was flashed around the circle. The Set annulus had an inner and outer radius of 0.7 and 0.75 deg and was flashed for 48 ms. Trials were aborted if the eye moved outside the fixation window or hand fixation was broken before Set.

For responses made after Set, the produced interval (*t*_*p*_) was measured from the endpoint of Set to the moment the saccade was initiated (eye trial) or the button was triggered (hand trial). Reward was provided if the animal used the correct effector and *t*_*p*_ was within an experimentally controlled acceptance window. For saccade responses, reward was not provided if the saccade endpoint was more than 2.5 deg away from the extinguished saccade target, or if the saccade endpoint was not acquired within 33 ms of exiting the fixation window.

The width of the reward acceptance window was adjusted adaptively on a trial-by-trial basis and independently for each condition using a one-up one-down staircase procedure. As such, animals received reward on nearly half of trials (57% in monkey A and 51% in monkey D), and the magnitude of the reward scaled linearly with accuracy. Additionally, rewarded trials were accompanied by brief visual and auditory feedback. For the visual feedback, either the color of the saccade target (for Eye trials) or the central square (for the Hand trials) turned green.

#### No-memory control task

To validate our hypothesis that slow fluctuations in animals’ behavior arose from memory fluctuations, we performed a control experiment in two naive monkeys. In the control experiment, the animals did not have to remember the target interval *t*_*t*_, but instead measured it on every trial. This was done by presenting an additional flash (“Ready”) shortly before the Set flash such that the interval between Ready and Set was fixed and equal to *t*_*t*_. This effectively removed the need for the animal to hold the target interval in memory. We limited the control experiment to a single effector (Eye) and a single interval (*t*_*t*_ = 840 ms).

#### Electrophysiology

Recording sessions began with an approximately 10-minute warm-up period to allow animals to recalibrate their timing and exhibit stable behavior. We recorded from 932 single- or multi-units in the thalamus, 568 units in the dorsomedial frontal cortex (DMFC), and 509 units in caudate, using 24-channel linear probes with 100 μm or 200 μm interelectrode spacing (V-probe, Plexon Inc.). The number of simultaneously recorded neurons across sessions were shown in Figure S11A (mean ± s.d.; Thalamus: 17.9 ± 9.0; DMFC: 9.2 ± 4.3; Caudate: 14.8 ± 6.5). The DFMC comprises supplementary eye field, dorsal supplementary motor area (i.e., excluding the medial bank), and pre-supplementary motor area. We recorded from the left hemisphere from Monkey A and right from Monkey D, which was contralateral to the preferred hand used in the button press. Recording locations were selected according to stereotaxic coordinates with reference to previous studies as well as each animal’s structural MRI scan. The region of interest targeted in the thalamus was within 1 mm of antidromically identified neurons. All behavioral and electrophysiological data were timestamped at 30 kHz and streamed to a data acquisition system (OpenEphys). Spiking data were bandpass filtered between 300 Hz to 7 kHz and spike waveforms were detected at a threshold that was typically set to 3 times the RMS noise. Single- and multi-units were sorted offline using custom software, MKsort (https://github.com/ripple-neuro/mksort).

#### Antidromic Stimulation

We used antidromic stimulation to localize DMFC-projecting thalamic neurons. Antidromic spikes were recorded in response to a single biphasic pulse of duration 0.2 ms (current < 500 uA) delivered to DMFC via low impedance tungsten microelectrodes (100 – 500 KΩ, Microprobes). A stainless steel cannula guiding the tungsten electrode was used as the return path for the stimulation current. Antidromic activation evoked spikes reliably at a latency ranging from 1.8 to 3ms, with less than 0.2 ms jitter.

### Human experiments

Each experimental session lasted approximately 60 minutes. Each subject completed 2-3 sessions per week. Similar to monkeys, experiments were conducted using the MWorks. All stimuli were presented on a black background monitor. Subjects were instructed to hold their gaze on a fixation point and hold a custom made push button using their dexterous hand, throughout the trial. Subjects viewed the stimuli binocularly from a distance of approximately 67cm on a 23-inch monitor (Apple, A1082 LCD) driven by a Mac Pro at a refresh rate of 60 Hz in a dark and quiet room. Eye positions were tracked with an infrared camera (Eyelink 1000 plus, SR Research Ltd.) and sampled at 1 kHz. The state of the button was converted and registered as digital TTL through a data acquisition card (National Instruments, USB-6212). The Cue-Set-Go task for humans was similar to monkeys with the following exceptions: (1) in each session, we used a single *t*_*t*_ sampled from a normal distribution (mean: 800ms, std: 80ms); (2) the saccadic target was 10 deg (instead of 8 deg) away from the fixation point. On average, 50.2% of trials received positive feedback.

A set of different human subjects performed the same timing interval production task, except that the nature of the feedback was provided probabilistically. Subjects received “correct” feedback with the probability of 0.7 when *t*_*p*_ was within a window around *t*_*t*_, and with the probability of 0.3 when errors were outside that window (Fig. 4C). The window length was adjusted based on the overall behavioral variability so that each subject receives approximately 50% “correct” feedback in each behavioral session. Only one effector - hand pressing - was used in this causal experiment, as we have established that both the slow and reward regulated variability were effector specific. All the experimental parameters were set identical to the main experiment.

## Data Analysis

All offline data processing and analyses were performed in MATLAB (2016b, MathWorks Inc.).

### Analysis of behavior

Behavioral data for the CSG task comprised of N = 203 behavioral sessions consisting of n = 167,115 trials in monkeys (N = 95, n = 71,053 for monkey A and N = 108, n = 96,062 for monkey D), N = 62 sessions and n = 59,297 trials in humans, and N = 51 sessions and n = 30,695 trials in the probabilistic feedback experiment in human subjects. Behavioral data for the no-memory control task was collected in N = 26 sessions consisting of n = 75,652 trials in two naive monkeys (N = 9, n = 32,041 for monkey G and N = 17, n = 43,611 for monkey H).

We computed the mean and standard deviation of *t*_*p*_, denoted by *μ*(*t*_*p*_) and *σ*(*t*_*p*_), respectively, for each trial type within each session (Figure 1C). We additionally analyzed local fluctuations of *μ*(*t*_*p*_) and *σ*(*t*_*p*_) by computing these statistics from running blocks of 50 trials within session and averaged across sessions. The resulting distribution of local *μ*(*t*_*p*_) and *σ*(*t*_*p*_) were shown in Figure S1. The mean of *σ*(*t*_*p*_) for each corresponding *μ*(*t*_*p*_) bin and the averaged reward across all trials in each *μ*(*t*_*p*_) bin were plotted in Figure 1D. Results were qualitatively unchanged when the block length was increased or decreased by a factor of two.

We also examined the slow fluctuations of *t*_*p*_ for pairs of trials that were either of the same type (e.g., Eye-Short versus Eye-Short) or of different types (e.g., Hand-Long versus Eye-Short). For trials of the same type, we computed partial correlation coefficients of *t*_*p*_ pairs by fitting a successive autoregressive model with the maximum order of 60 trial lag (Box *et al.*, 2015) (Figure 2A). 1% and 99% confidence bounds were estimated at 2.5 times the standard deviation of the null distribution. For trials of different types, we calculated the Pearson correlation coefficient of pairs of *t*_*p*_ of various lags (Figure 2B, Figure S2 and S6). To clarify our analysis, we use an example of how we estimated the cross correlation between pairs of HS - ES with a trial lag of 10: (1) normalize (z-score) two *t*_*p*_ vectors associated with HS and ES in each session; (2) take pairs of HS-ES that are 10 trials apart within each session; (3) combine the pairs across sessions; (4) compute Pearson correlation coefficient. We also computed a corresponding null distribution from 100 randomly shuffled trial identity. 1% and 99% confidence intervals were estimated from the null distribution.

Finally, we quantified the mean and standard deviation of the relative error denoted by *μ*(*e*^*n*^) and *σ*(*e*^*n*^) as a function of error in the previous trial (*e*^*n-1*^) for each pair of trial types (Figure S4 A and B). Deriving reliable estimates of *μ*(*e*^*n*^) and *σ*(*e*^*n*^) as a function of *e*^*n-1*^ from a non-stationary process requires a large number of trials. Since each trial can be of four different types (ES, EL, HS, HL), consecutive trials comprise 16 distinct conditions (e.g., ES-EH, HL-EL, etc, Figure S4A shows the distribution for all conditions). The limited number of trials in each session limited the reliability of statistics estimated for each individual condition. To gain more statistical power, we combined results across trial types in two ways. First, for each effector, we combined the Short and Long trial types by normalizing *t*_*p*_ values by their respective *t*_*t*_. The resulting variable was defined as relative error *e*^*n*^ = (*t*_*p*_^*n*^-*t*_*t*_)/*t*_*t*_. This reduced the number of conditions by a factor of four, leaving consecutive trials that were either associated with the same effector or with different effectors (e.g., E-E, E-H, H-E, and H-H). We further combined trials to create a “same effector” condition that combined E-E with H-H, and a “different effector” condition that combined E-H with H-E. Animals and human subjects were allowed to take breaks during the experimental sessions. However, the pairs of consecutive trials used in all analyses, regardless of the trial condition, were restricted to the two consequent and completed trials that were no more than 7 seconds apart.

We examined the relationship between *μ*(*e*^*n*^) and *e*^*n-1*^ using a linear regression model of the form *μ*(*e*^*n*^) *= m*_*0*_*+m*_*1*_*e*^*n-1*^, and the relationship between *σ*(*e*^*n*^) and *e*^*n-1*^ using a quadratic regression model of the form *σ*(*e*^*n*^) *= s*_*0*_*+s*_*1*_*e*^*n-1*^*+s*_*2*_(*e*^*n-1*^)^*2*^. We assessed the significance of the monotonic relationship between *μ*(*e*^*n*^) and *e*^*n-1*^ by the slope of linear regression (*m*_*1*_), and the significance of the U-shaped profile in *σ*(*e*^*n*^) by the square term of a quadratic regression (*s*_*2*_). Note that the choice of these models was not theoretically motivated; we simply considered these regression models to be an approximate function for testing the profile of *μ*(*e*^*n*^) and *σ*(*e*^*n*^) as a function of *e*^*n-1*^. The 1% and 99% confidence intervals were defined as, e.g. *s*_*2*_ ± *t*_*(.01, N-p)*_ *SE(s*_*2*_), where s_2_ is the estimated coefficient, *t*_*(.01, N-p)*_ is the 1 percentile of t-distribution with N-p degrees of freedoms, and SE*(s*_*2*_) is the standard error of estimated *s*_*2*_.

We applied the same analysis to the probabilistic reward experiment in humans after grouping the trials based on whether the feedback in the previous trials was “correct” or “incorrect”. Before analyzing the variability, we removed outlier trials defined as trials in which *t*_*p*_ was more than three standard deviations away from the mean. Since subjects were allowed to initiate the trials, we also excluded pairs of adjacent trials that were more than 7 seconds apart. When combining data across sessions, we normalized the relative error across sessions and subjects. To avoid sampling bias, the same number of trials were drawn repeatedly after “correct” or “incorrect”. The standard error of the mean was computed from 100 repeats with replacement (Fig. 4D and S5C).

### Reward-sensitive Gaussian process (RSGP) model simulation and fitting

We constructed a reward-sensitive Gaussian process model whose covariance function, ***K***_*RSGP*_, is a weighted sum of two kernels, a traditional squared exponential kernel, for which we used subscript SE (***K***_*SE*_), and a reward-sensitive kernel with subscript RS (***K***_*RS*_). The two kernels contribute to *K*_*RSGP*_ through scale factors *σ*^*2*^_*SE*_ and *σ*^*2*^_*RS*_, respectively. In both kernels, the covariance term between any two trials (trial *n* and *n-r*) drops exponentially as a function of trial lag (*r*). The rates of drop for ***K***_*SE*_ and ***K***_*RS*_ are specified by characteristic length parameters, *l*_*SE*_ and *l*_*RS*_, respectively. The model also includes a static source of variance, *σ*^*2*^_*0*_*I* (***I*** stands for the identity matrix):

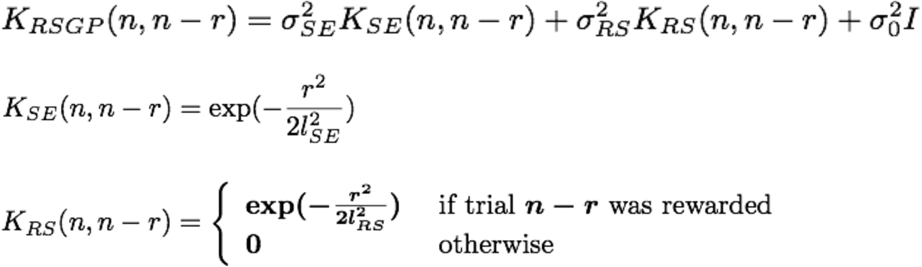

Note that ***K***_*RS*_ values depend on reward history and are thus not necessarily invariant with respect to time; i.e. *K*(*n, n* − *i*) ≠ *K*(*m, m* − *i*). This formulation allows past rewarded trials to have higher leverage on future trials and this effect drops exponentially for rewarded trials farther in the past.

We simulated the RSGP by applying GP regression based on the designated covariance function (Table 3). To simplify our formulations and without loss of generality, we replaced *σ*^*2*^_*SE*_ and *σ*^*2*^_*RS*_ by *σ*^*2*^ and (1 -)*σ*^*2*^, respectively where = 1.0, 0, and 0.5 for the three examples (Figure 3A, Table S1).

**Table 3.**
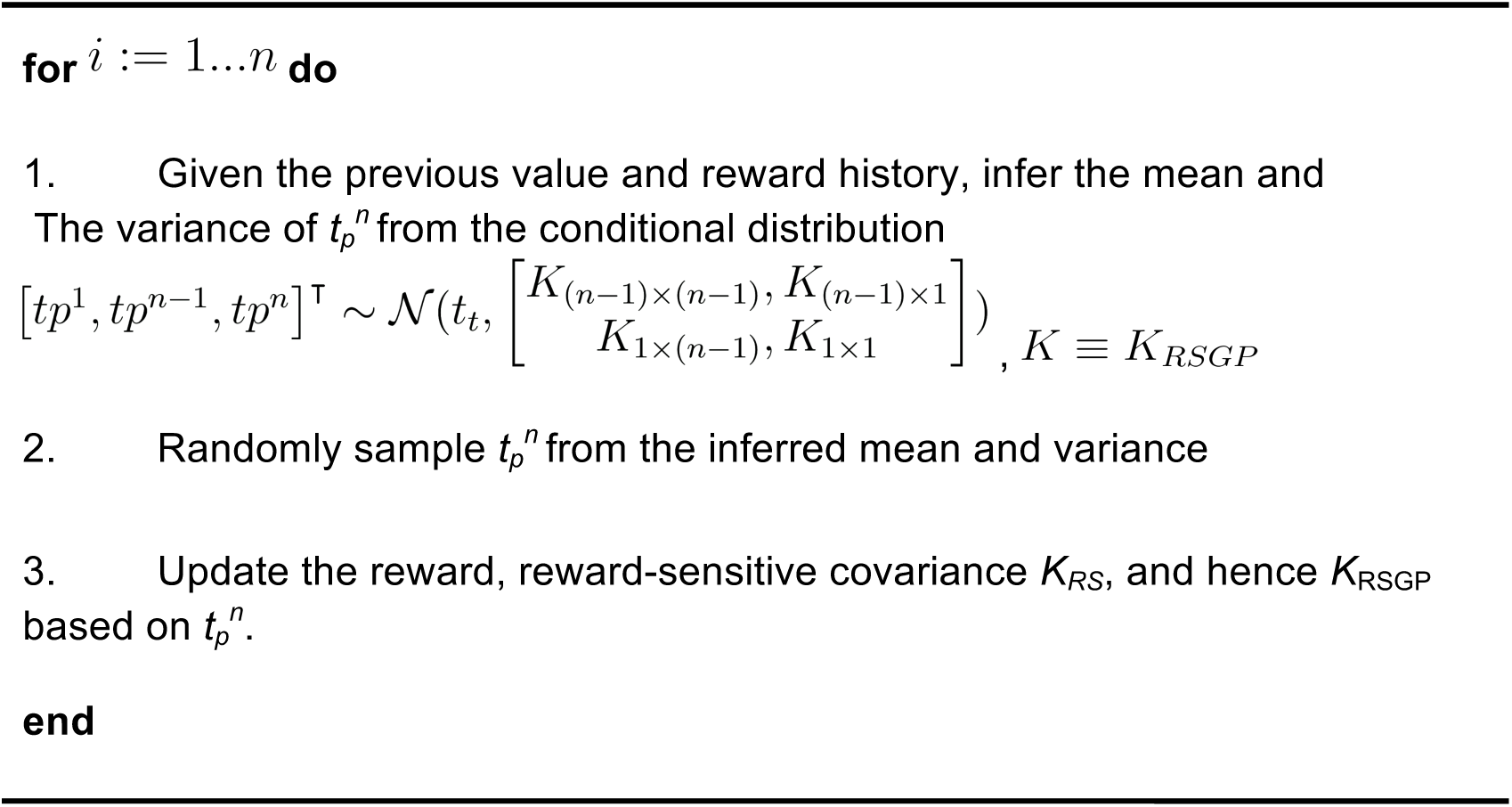
Algorithm for generating time series based on RSGP model.

As both the slow fluctuation and reward regulation were context specific, we fit the model to behavioral data for each trial type (ES, EL, HS, and HL) separately. To do so, we ordered *t*_*p*_ values associated with the same trial type within each behavioral session chronologically and treated them as consecutive samples from the model irrespective of the actual trial lag between them. Although this strategy made the model fitting more tractable, the inferred length constants in units of trials are likely smaller than the true values in the data. For the same reason, the temporal distance in the kernel function was different from the actual trial lag (Figure 2B, S7). Methods for fitting behavioral data to the RSGP model were adapted from *Gaussian Processes for Machine Learning* (Rasmussen and Williams, 2006). The objective was to maximize the marginal likelihood of the observed data with respect to the hyperparameters {*l*_*SE*_, *σ*_*SE*_, *l*_*RS*_, *σ*_*RS*_, *σ*_*0*_}. Using simulations, we found that optimization through searching the entire parameter space was inefficient and hindered convergence. Therefore, we implemented a two-step optimization. We first used the unrewarded trials to estimate *l*_*SE*_ and *σ*^*2*^_*SE*_, and then used those fits to search for the best fit of the remaining hyperparameters (*l*_*RS*_, *σ*_*RS*_, and *σ*_*0*_) using all trials. The optimization of the multivariate likelihood function was achieved by line searching with quadratic and cubic polynomial approximations. The conjugate gradients was used to compute the search directions (Rasmussen and Williams, 2006). The landscape of likelihood indicated that the optimization was convex for a wide range of initial values (Figure S6A, Table S1).

The RSGP model fit to data provides a prediction of the distribution of *t*_*p*_ on each trial based on previous trials (*t*_*p*_ and reward history). We used this distribution to generate simulated values of *t*_*p*_ for each session, and repeated this process (n = 100) to estimate the distribution of *μ*(*e*^*n*^) and *σ*(*e*^*n*^) in relation to *e*^*n-1*^ using the same analysis we applied to behavioral data (Figure 5C). To derive an estimate of the slow part of error, *e*_slow_, we first fitted the RSGP to the behavior, and then used the reduced RSGP that only included the slow kernel (***K***_*SE*_) to predict the expected value of *e*_slow_, i.e. the mean of a GP process governed by ***K***_*SE*_.

### RSGP model for speed

We reformulated RSGP such that the key variable in the model was speed – not the interval (Figure S9 A). Accordingly, the speed signal was used to control the slope of a ramping activity that triggered an action when it breached a fixed threshold. The minor modification from target interval to speed enabled the RSGP to capture both the dependence of variability on reward within each condition and the scalar variability across conditions (Figure S9 B,C).

### Markov chain Monte Carlo (MCMC) model

The MCMC model was proposed by Haith and Krakauer (Haith and Krakauer, 2014). We adapted this algorithm to our task by adding time production noise to step 3, and approximating the cost as the opposite of reward (i.e. *J*(*t*_*p*_) = - *V*(*t*_*p*_)). and simulated it to test whether it could capture (1) the long-term correlations, (2) the monotonic relationship between the mean of error on trial n, *μ(e*^*n*^), and error on trial n-1, *e*^*n-1*^, and (3) the U-shaped relationship between the standard deviation of error, *σ(e*^*n*^), and *e*^*n-1*^. If we denote the variable of interest, *t*_*p*_, and the reward associated with it, V(*t*_*p*_), the algorithm they proposed works as follows:

**Table 4.**
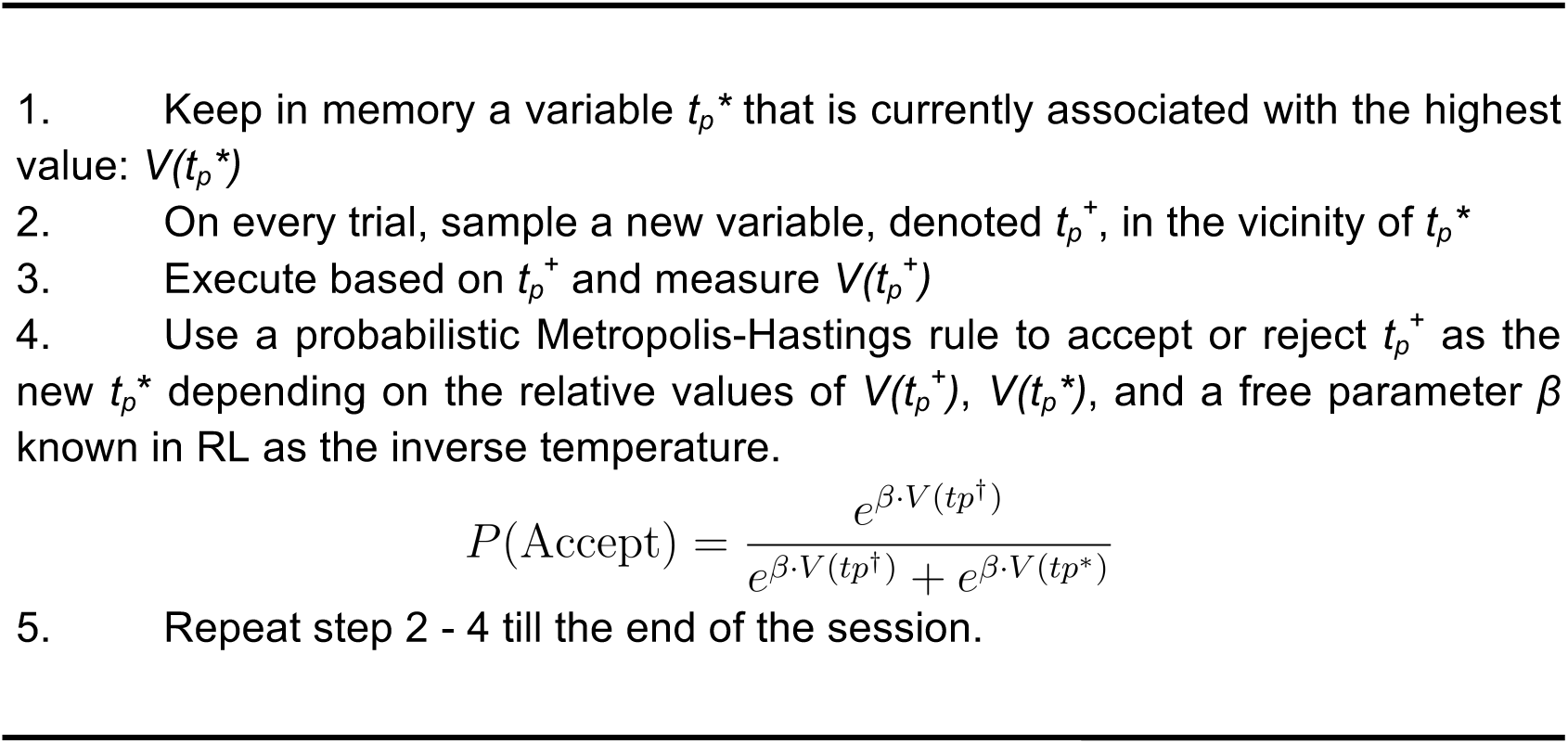
Algorithm for generating time series based on MCMC.

We have extensively explored the parameter space of this model. In the simulation (Figure S5 A), we used 0.1\**t*_*t*_ and 0.05\**t*_*t*_ as the level of sampling noise in step 2, and execution noise in step 3, respectively. The inverse temperature *β* was 100.

### Relationship between neural activity and the slow component of behavior

We used linear regression to examine whether and to what extent the population neural activity could predict the slow component of error (*e*_slow_) inferred from the RSGP model fits to behavior (as described in the previous section) using the following regression model:

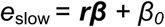

where ***r*** represents a matrix (*nxN*) containing spike counts within a 250 ms of *N* simultaneously recorded neurons across *n* trials, *β*_0_ is a constant, and ***β*** is an *N*-dimensional vector specifying the contribution of each neuron to *e*_slow_. We used a random half of trials (training dataset) to find ***β*** and *β*_*0*_ and the other half (validation dataset) to test the model, and quantify the success of the model by computing the Pearson correlation coefficient (Figure 6B-D) between the *e*_slow_ inferred from RSGP model fits to behavior and *e*_slow_ predicted from the neural data using the regression model.

We initially tested the regression model using spike counts immediately before Set and later extended the analysis to different time points throughout the trial. To do so, we aligned spike times to various events throughout the trial (Cue, Tar, Set, Go) and tested the regression model every 125 ms around each event (4 time points after Cue, 3 time points before Tar, 3 time points after Tar, 4 time points before Set, 4 time points after Set and 4 time points before Go).

To ensure that the model was predictive and not simply overfitting noisy spike counts, we used a cross-validation procedure: for each session, we used a random half of the trials to estimate ***β***_Th_, and the other half to quantify the extent to which *z*_Th_ could predict *e*_slow_. Note that some correlations are negative because of cross-validation (we used a random half of data to estimate the drift direction and the other half for estimation correlations). Otherwise, all correlations should have been non-negative. Note that the number of sessions was combined across all 4 trial types as shown in Figure 6B-D bottom.

### Statistical Analysis

Mean ± standard deviation (SD), Mean ± standard error of the mean (SEM) or median ± median absolute deviation (MAD) were used to report statistics. We detailed all the statistics in the Results and figure captions. All hypotheses were tested at a significance level of 0.01 and P-values were reported. We used t-tests to perform statistical tests on the following variables: (1) weber fraction (ratio of standard deviation to mean of *t*_*p*_), (2) cross correlation between pairs of trials of different lag and trial type, (3) modulation of the variability by discrete or graded reward (4) variance terms in the RSGP model (*σ*^*2*^_*SE*_, *σ*^*2*^_*RS*_, and *σ*^*2*^_*0*_) which were assumed to be normally distributed. We used one-tailed paired, two-tailed paired or two-sample *t-*tests depending on the nature of data and question. The length scale parameters of the RSGP model (*l*_*SE*_ and *l*_*RS*_) were not normally distributed. Therefore, we used a one-way ANOVA to test whether the two were significantly different. We used a two-sample *F*-test to compare the variability of production interval for different pair-trial conditions (H0: equal variance).

### Data Availability Statement and Accession Code Availability Statements

Data that support the findings of this study are available from the corresponding authors. Matlab codes for RSGP simulation and model fitting are available at https://github.com/wangjing0/RSGP.

### Mathematical notation

| Symbol | Description |
| --- | --- |
| $t_p^n$ | Production time in the $n^{\text{th}}$ trial |
| $e^n$ | $(t_p^n - t_t)/t_t$, relative error of production time in the $n^{\text{th}}$ trial |
| $\mu(e)$ | Mean of relative error |
| $\sigma(e)$ | Standard deviation of relative error |
| $K(i, j)$ | Covariance between trial $i$ and trial $j$ |
| $\sigma^2$ | Signal variance or noise variance associated with a Gaussian process |
| $l$ | The length scale of the squared-exponential covariance function |
| $\mathbf{r}^n(t)$ | Population spiking at time $t$ in the $n^{\text{th}}$ trial |
| $\beta$ | Regression coefficient |
| $z$ | Projection of population spiking activity onto a low dimensional representation |

## Author Contributions

J.W., E.H. and M.J. conceived the project. J.W. and E.H. collected the main behavioral and electrophysiology data. J.W. analyzed the data and developed the model. E.H. played a major role in data analysis. J.W. and M.J. interpreted the data with contribution from E.H. and N.M. N.M. performed and analyzed the control experiment in monkeys. J.W. designed and analyzed the human psychophysics experiments and A.A. conducted the experiments. M.J. supervised the project. All authors contributed to the writing of the manuscript.

## Acknowledgments

M.J. is supported by NIH (NINDS-NS078127), the Sloan Foundation, the Klingenstein Foundation, the Simons Foundation, the McKnight Foundation, and the McGovern Institute. N.M. is supported by the Center for Sensorimotor Neural Engineering.

## Data and code availability

The data and custom analysis scripts that support the findings of this study will be made available publicly on a designated website immediately after publication.

## Supplementary information

**Figure S1.**
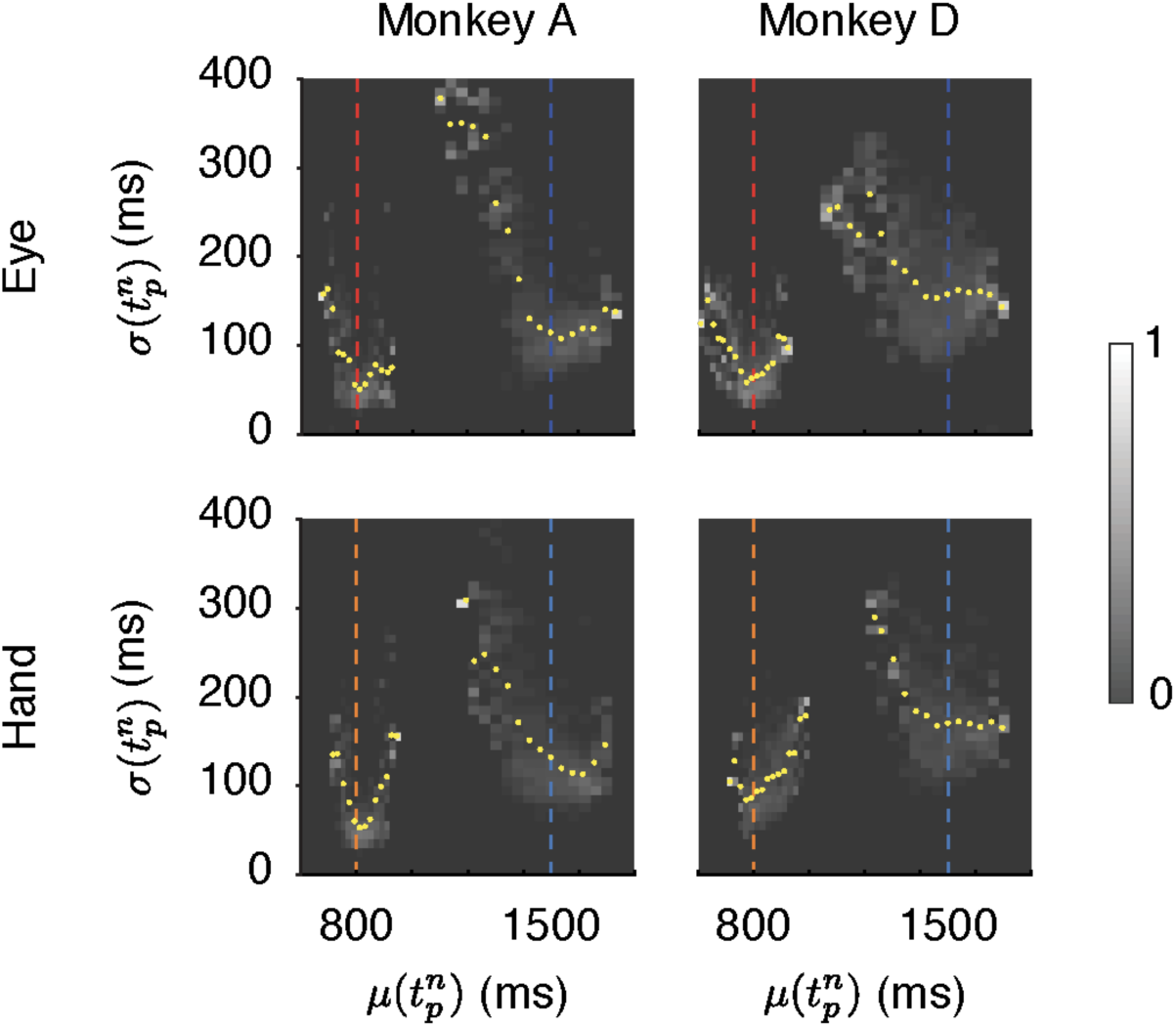
Dependence of timing variability on reward rate. The gray scale shows the marginal distribution of *σ*(*t*_*p*_) for given *μ*(*t*_*p*_) derived from local estimates of *σ*(*t*_*p*_) and *μ*(*t*_*p*_) using a 50-trial sliding window (yellow dots: mean of *σ*(*t*_*p*_) for each *μ*(*t*_*p*_) value; colored dash lines: target intervals centered at 800 and 1500 ms). Bin size was 20 ms.

**Figure S2.**
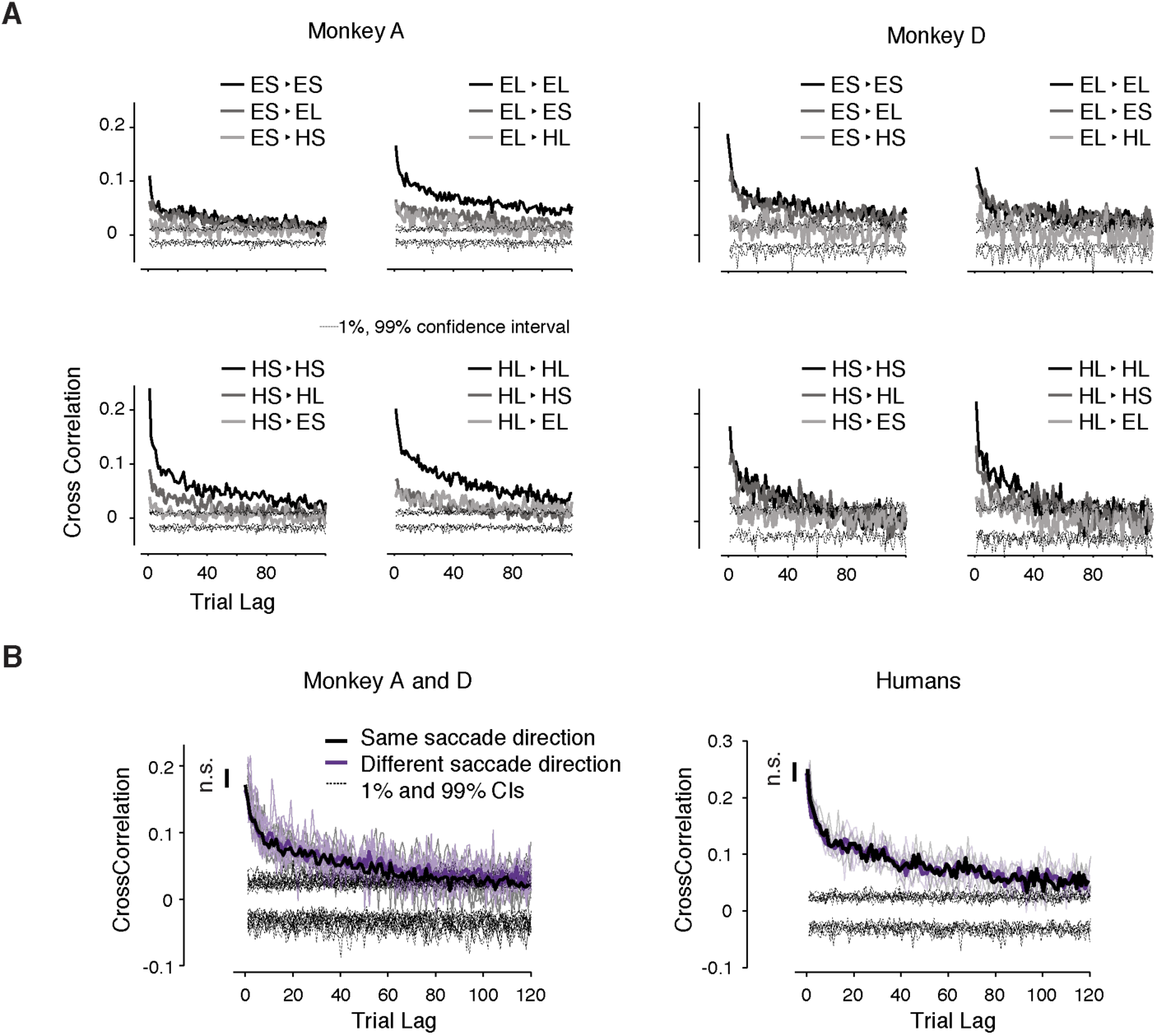
Context-dependent slow fluctuations of timing variability for (A) transitions between different trial types and for (B) different saccade directions. Same as Figure 2B shown for the two animals and different trial types separately. **(B)** Similar to Figure 2B, it shows the long range correlation between trials with same and different saccade direction separately. The analysis was performed separately for trials of the same type (i.e., same target interval and same effector), and pooled afterward (black: same direction; purple: different direction; bold: average; dashed [1%-99%] confidence intervals). The long-term correlations were not significantly different for the same and different directions (Monkeys: paired sample t-test, p = 0.39, t = 0.86, df = 159. Humans: paired sample t-test, p = 0.55, t = 0.60, df = 79).

**Figure S3.**
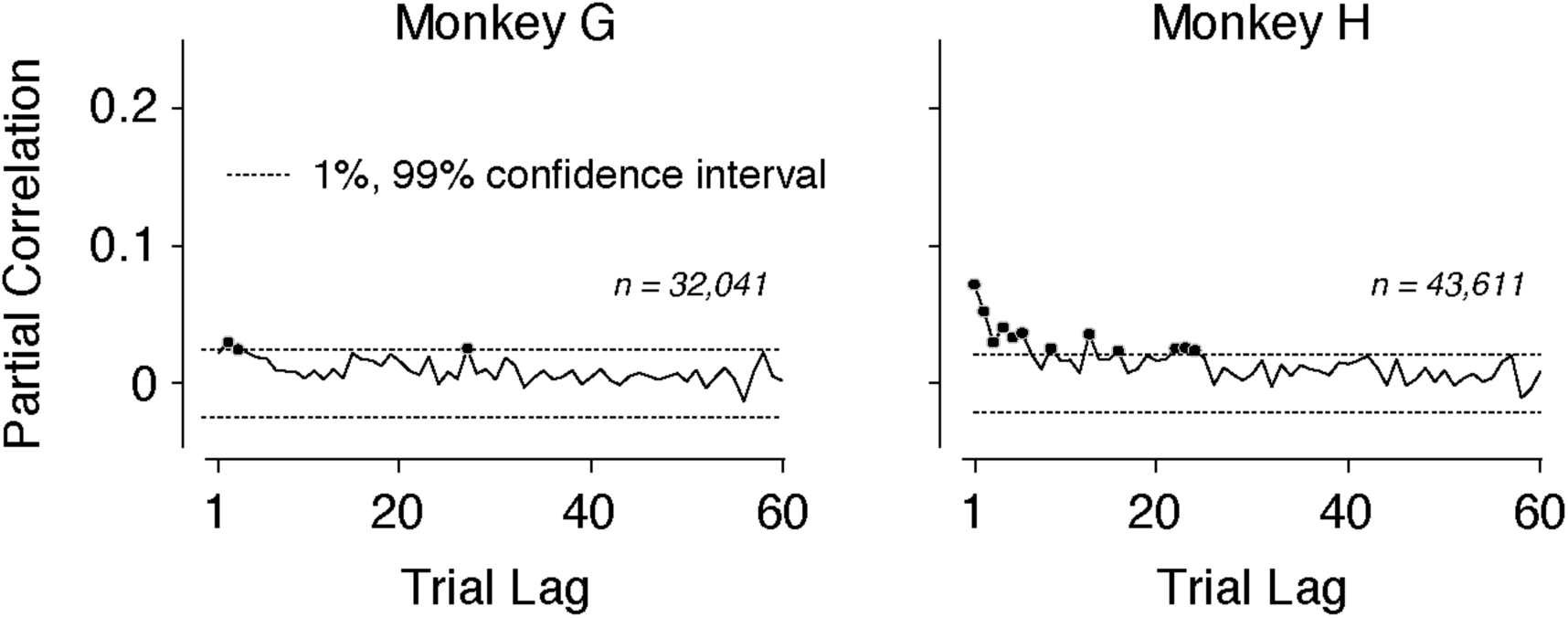
Serial correlation of *t*_*p*_ in the control experiment. Slow fluctuations of *t*_*p*_ were diminished in the control experiment in which the demand for remembering the target interval was minimized. Results are shown in the same format as in Figure 2A.

**Figure S4.**
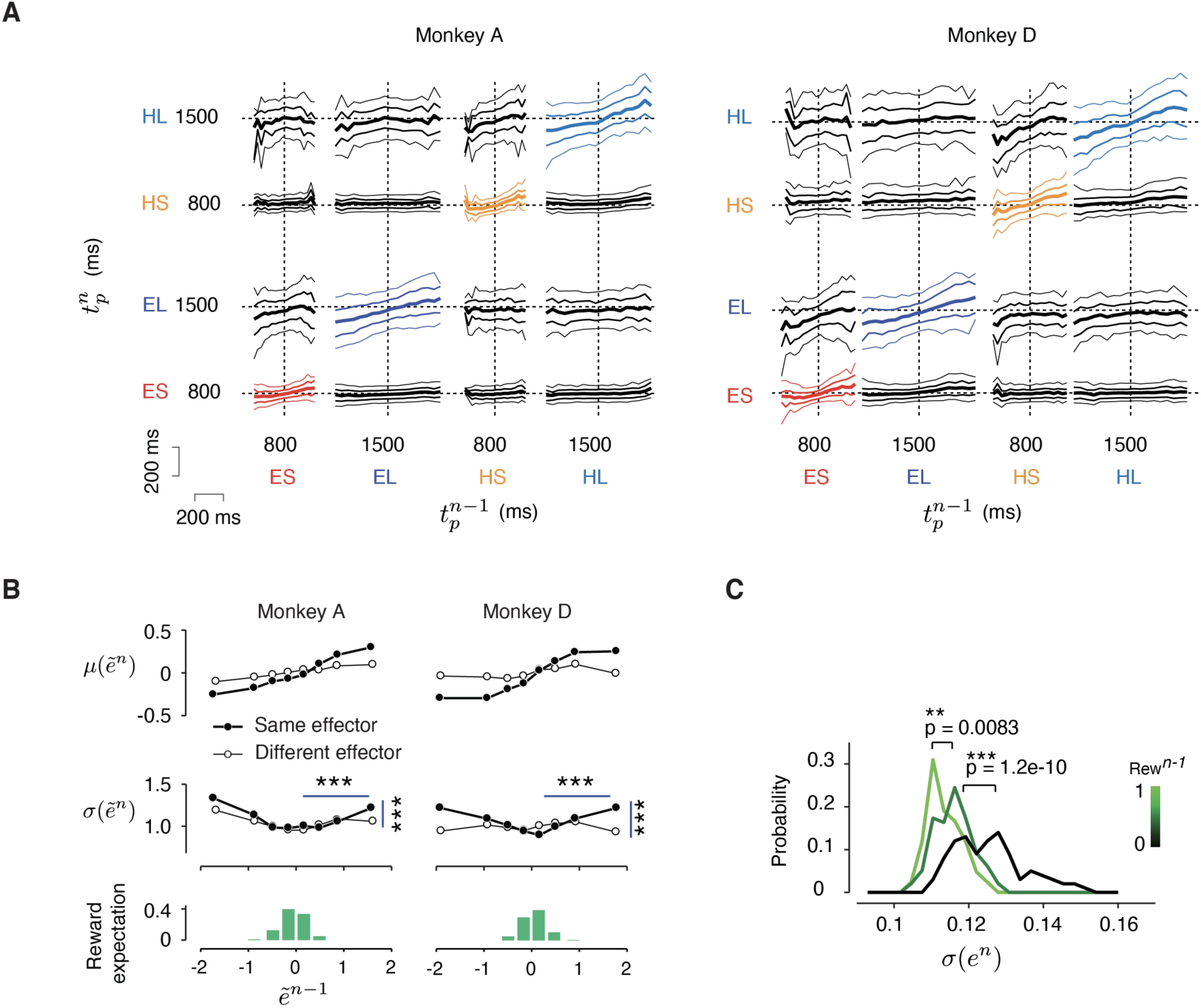
The effect of reward on behavioral variability. **(A)** Cross-condition relationship between produced interval (*t*_*p*_) in consecutive trials. The results for each animal are organized on a 4×4 panel covering the full set of 16 possible transitions between 4 trial types (ES: Eye-Short; EL: Eye-Long; HS: Hand-Short; HL: Hand-Long) with the abscissa and ordinate showing *t*_*p*_ at trial *n-1* (*t*_*p*_^*n-1*^) and n (*t*_*p*_^*n*^), respectively. In each panel, the median *t*_*p*_^*n*^ as the function of *t*_*p*_^*n-1*^ is marked in bold line at the center. The 25^th^ and 75^th^ percentile are marked with medium thickness, and the 10^th^ and 90^th^ percentile with thin lines. Bin size was 20 ms. **(B)** Error statistics within each session. Since our original analysis was based on data combined across sessions, one potential concern we had was that modulation of *t*_*p*_ variance with reward was due to differences across (but not within) sessions. To address this, we first removed outlier *t*_*p*_ values (more than 3 s.d. away from the mean) and then analyzed z-scored errors within each session (denoted 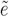). This normalization ensured that error pairs in every session were drawn from a zero-mean and unit-variance distribution. Results are shown in the same format as in Figure 3B and C. The variability increased significantly after unrewarded trials compared to that after rewarded trials (horizontal lines, *** p < 0.001; monkey A: F(8889,10621) = 1.15 for negative 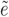 and F(9841,11103) = 1.09 for positive 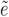; monkey D: F(21216,12364) = 1.25 for negative 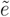 and F(1600,19293) = 1.38 for positive 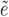), and this effect was context specific (vertical lines, for monkey A: F(10621,9841) = 1.10 and monkey D: F(12364, 16100) = 1.11). **(C)** Behavioral variability as a function of the magnitude of reward in the preceding trial. The distribution of *σ(e*^*n*^) sorted into three bins depending on reward magnitude (black: no reward; dark green: low reward; light green: high reward). The variability was significantly larger in the absence of reward in comparison to low reward (***, p << .001, two-sample t-test, t = 6.6, df = 117), and significantly larger in the low compared to the high reward (**, p = 0.008, two-sample t-test, t = 2.5, df = 68). Results were generated using bootstrapping; i.e., we computed *σ(e*^*n*^) from randomly sampled trials without replacement within each bin of *e*^*n-1*^.

**Figure S5.**
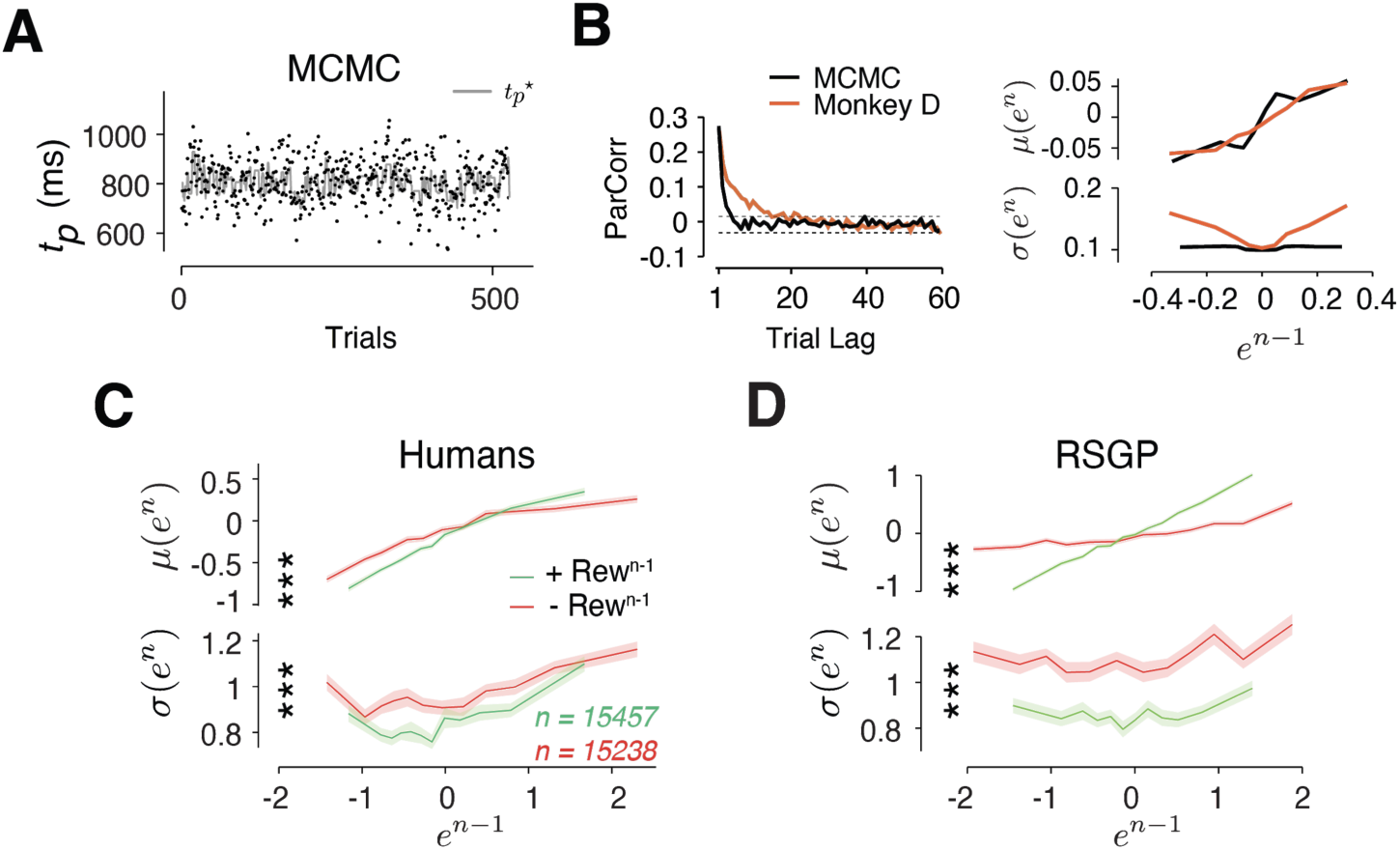
Analysis of error statistics for alternative generative processes and the causal experiment. **(A)** A Markovian sampling model. Production interval (*t*_*p*_) was generated as samples from a Markov chain Monte Carlo (MCMC) model (see Methods). **(B)** This model does not exhibit the long-term autocorrelations observed in animals’ behavior. Moreover, when the model was parameterized to capture the monotonic relationship between *μ(e*^*n*^) and *e*^*n-1*^, it failed to capture the U-shaped relationship between *σ(e*^*n*^) and *e*^*n-1*^ (*s*_*2*_ = 0.045 [-0.048 0.14]). Example data from Monkey D is shown in orange. **(C)** The effect of reward on behavioral variability in the probabilistic reward experiment. Result was combined for all subjects in Fig. 4D and was plotted in the same format as Fig. 3B. To avoid sampling bias, same number of trials were drawn repeatedly after “incorrect” (red, –*Rew*^*n-1*^) or “correct” (green, +*Rew*^*n-1*^) trials (Shaded area: standard error of the mean computed from 100 bootstraps). Behavioral variability *σ*(*e*^*n*^) increased significantly after ‘incorrect’ trial outcome regardless of the size of error (*** p<0.001, two-sample t-test on *σ*(*e*^*n*^) between the two conditions, t_198_ > 6.70 for all bins). The slope of the *μ*(*e*^*n*^) as a function of *e*^*n-1*^ was larger after ‘correct’ outcome (*** p<0.001, two sample t-test t_198_ = 16.8) demonstrating stronger behavioral correlation after positive reinforcement. **(D)** Same as (C) for the behavior of the RSGP model fit to the subject’s behavior.

**Figure S6.**
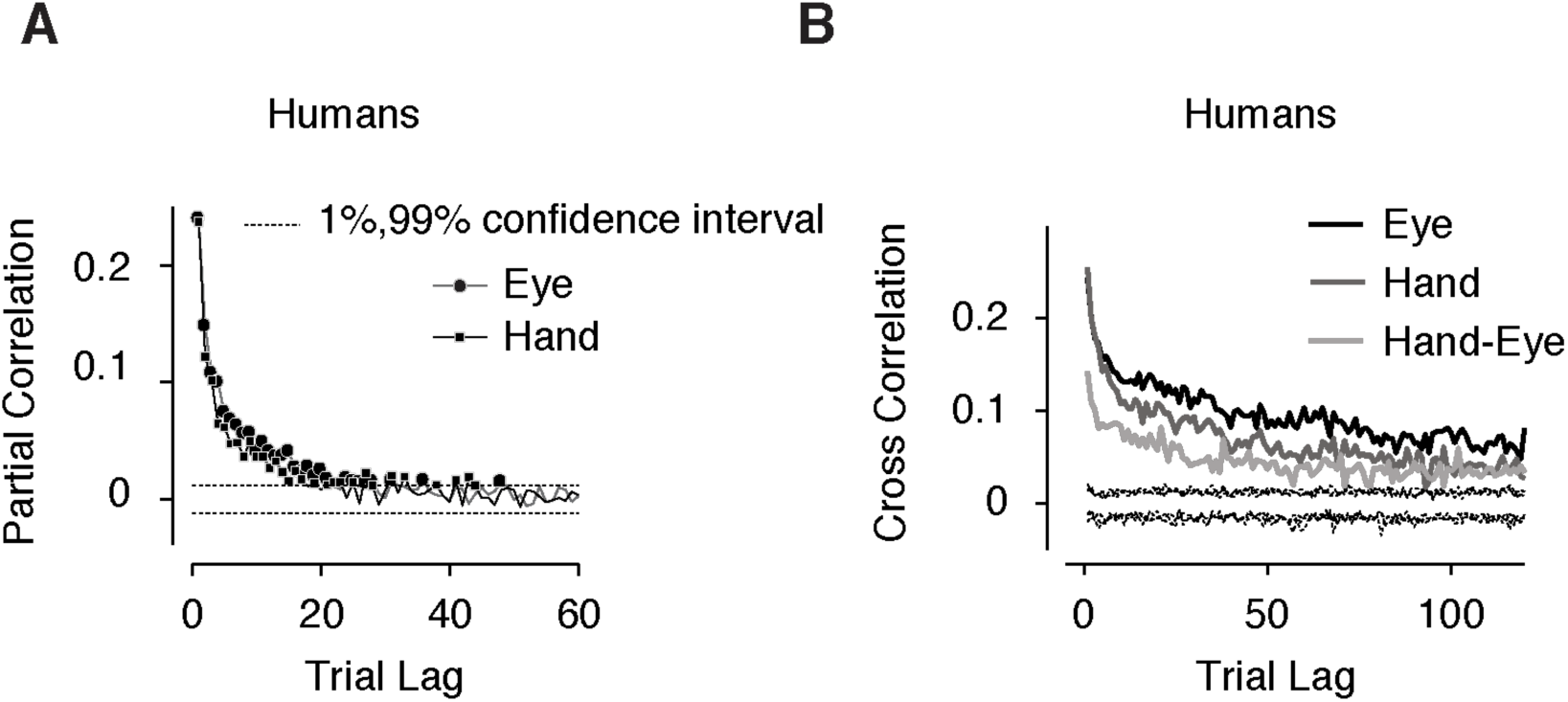
Long-term correlation of *t*_*p*_ and context-specificity in humans. Correlation was averaged across human subjects (N = 5). Same format as in Figure 2A and B.

**Figure S7.**
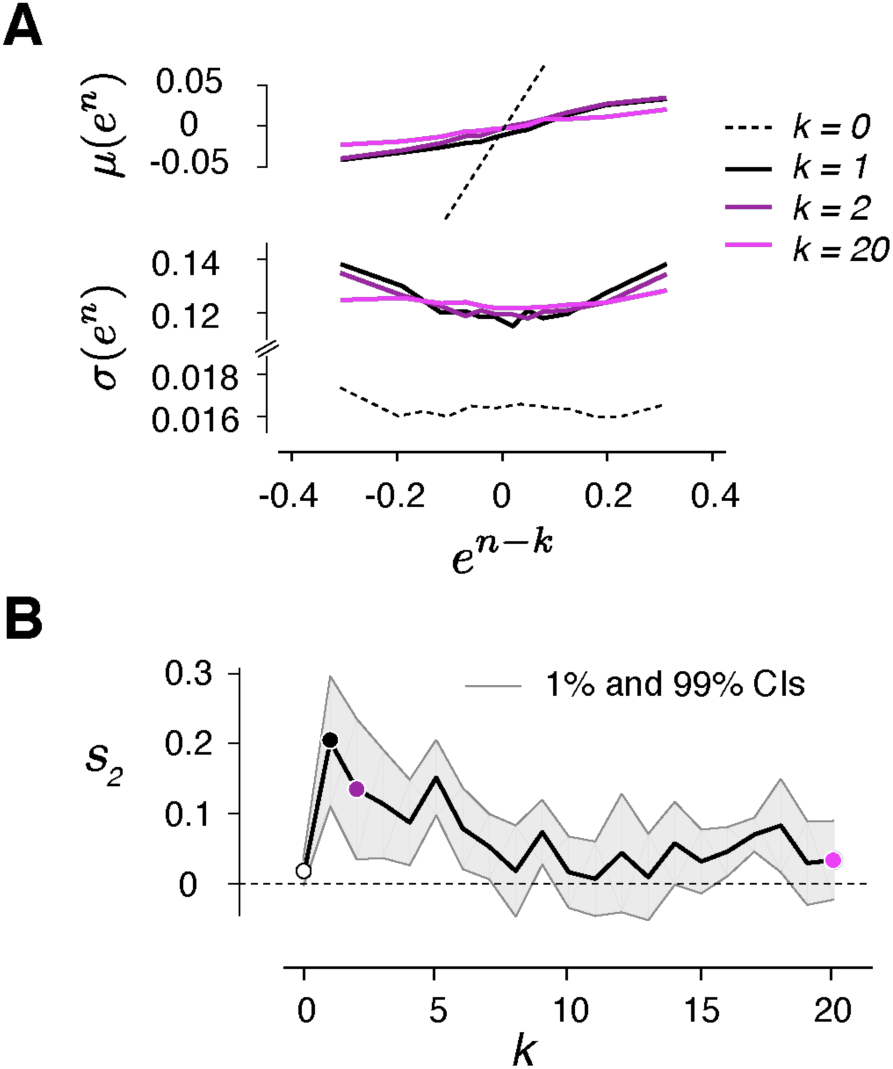
Error statistics as a function of trial lag. **(A)** Top: Average error on trial n, *μ(e*^*n*^) as a function of error in trial *n-k* (*e*^*n-k*^) for k = 0, 1, 2, and 20 (different colors). Bottom: Standard deviation of error on trial n, *σ(e*^*n*^) as a function of *e*^*n-k*^. Results were computed for each trial type separately averaged afterwards. **(B)** Modulation of variability by reward as a function of trial lag. Similar to the main paper, we tested the U-shaped profile using quadratic regression. The ordinate shows the coefficient of the square term (*s*_*2*_) of the quadratic regression as a function of trial lag (circles: *s*_*2*_ for the k values in A; shaded area: [1%-99%] confidence intervals). The reward has no direct bearing on the variability of the current trial (*k=0*), and its effect drops as a function of trial lag.

**Figure S8.**
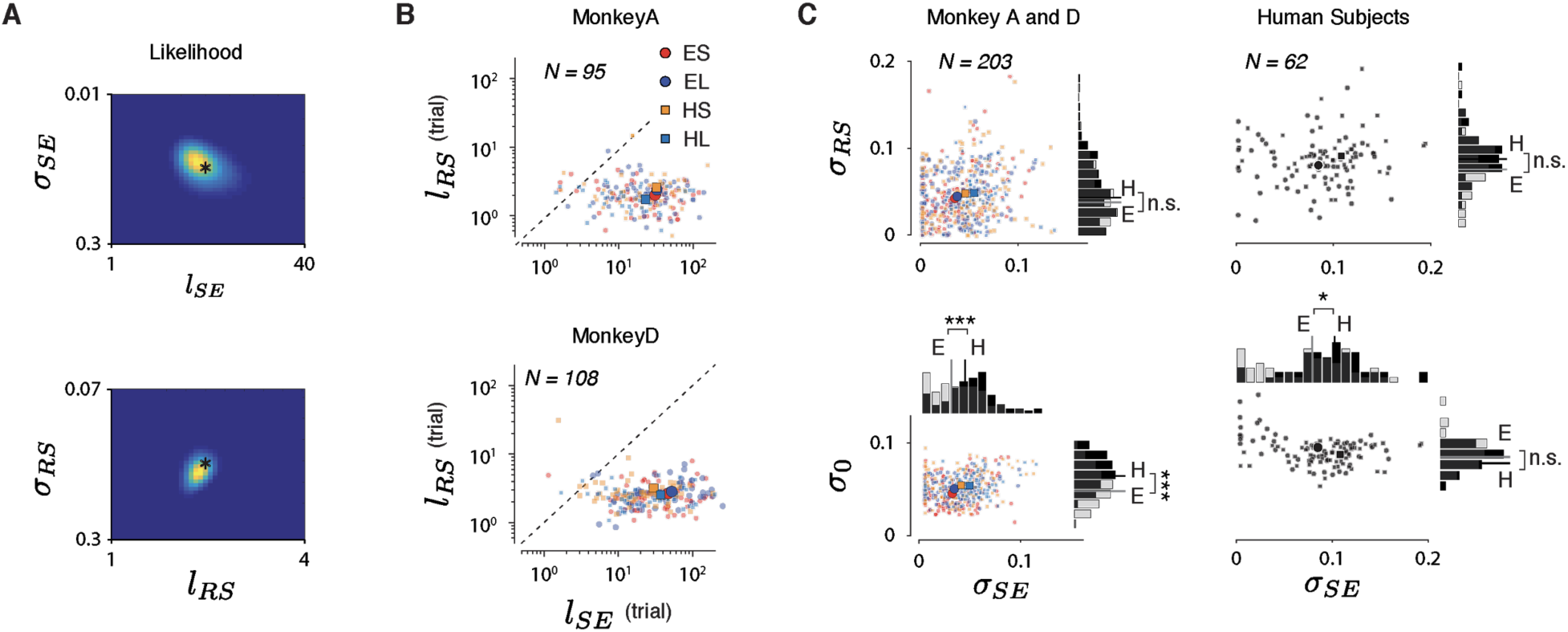
Results of the RSGP model fit to the behavior. **(A)** Simulation results indicated that the likelihood function captures the ground truth and that the error surface is convex for a wide range of initial values. See Methods for details. The hyperparameters used for simulation are marked by asterisks and the results of fitting are in Table S1. **(B)** Same as Figure 5B shown for the two animals separately. **(C)** The model fit of variances (*σ*_SE_, *σ*_RS_ and *σ*_0_) across sessions for monkeys (left column) and humans (right column).

**Figure S9.**
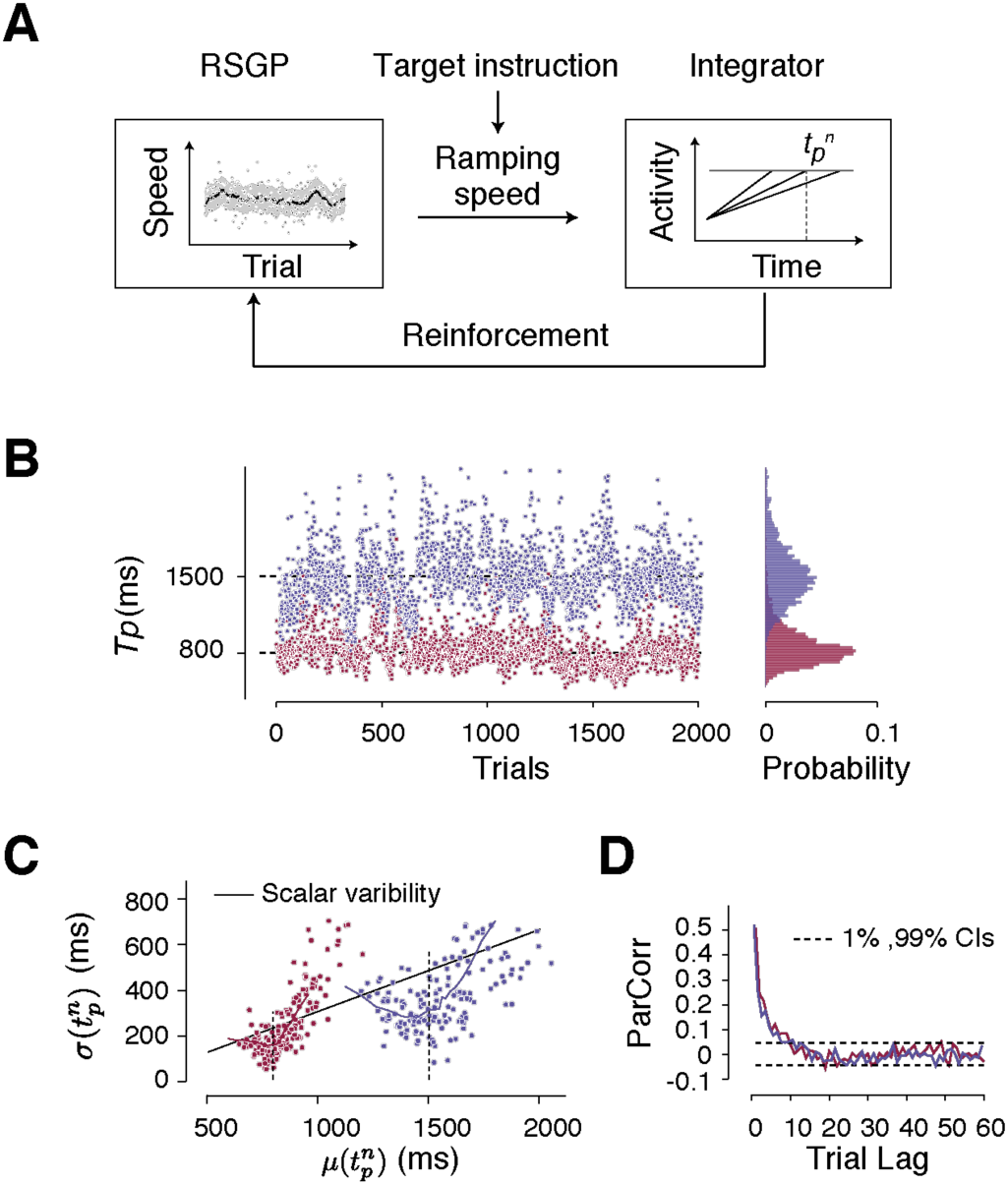
RSGP model for speed. **(A)** The model has two modules. The first module provides a tonic speed input, and the second module integrates the speed and generates a ramp toward a fixed (i.e. interval-independent) action-triggering threshold. The speed controls the slope of the ramp and therefore, the resulting produced interval (*t*_*p*_). Similar to the task, the presence or absence of reward depends on the error in *t*_*p*_ relative to the desired interval. The RSGP controls the output of the first module, and governs both the slow fluctuations of the speed (i.e., drift across trials), and the sensitivity of the speed to previous trials’ reward (reinforcement). Parameters used for generating speed were identical to those used previously in the RSGP (Figure 5A, Table S1). The only difference is that the model incorporates the instruction about the target interval, *t*_*t*_, by applying a gain factor to the output of the first module that is inversely proportiosnal to *t*_*t*_. This ensures that, on average, the ramp would reach the threshold at the instructed time. **(B)** *t*_*p*_ distribution of the model in an example session for the Short and Long conditions. **(C)** The relationship between the mean and standard deviation of *t*_*p*_ based on simulations of the model. The results were produced using the same analysis used in Figure 1D. The model captures the increase of variability for the Long relative to Short condition (i.e., scalar variability) as evident from the slope of the regression line (solid line; positive slope, p < 0.01). It also captures the U-shaped sensitivity of variability to reward around the desired interval (dashed line) for each condition. **(D)** Partial correlation of *t*_*p*_ shows slow fluctuations.

**Figure S10.**
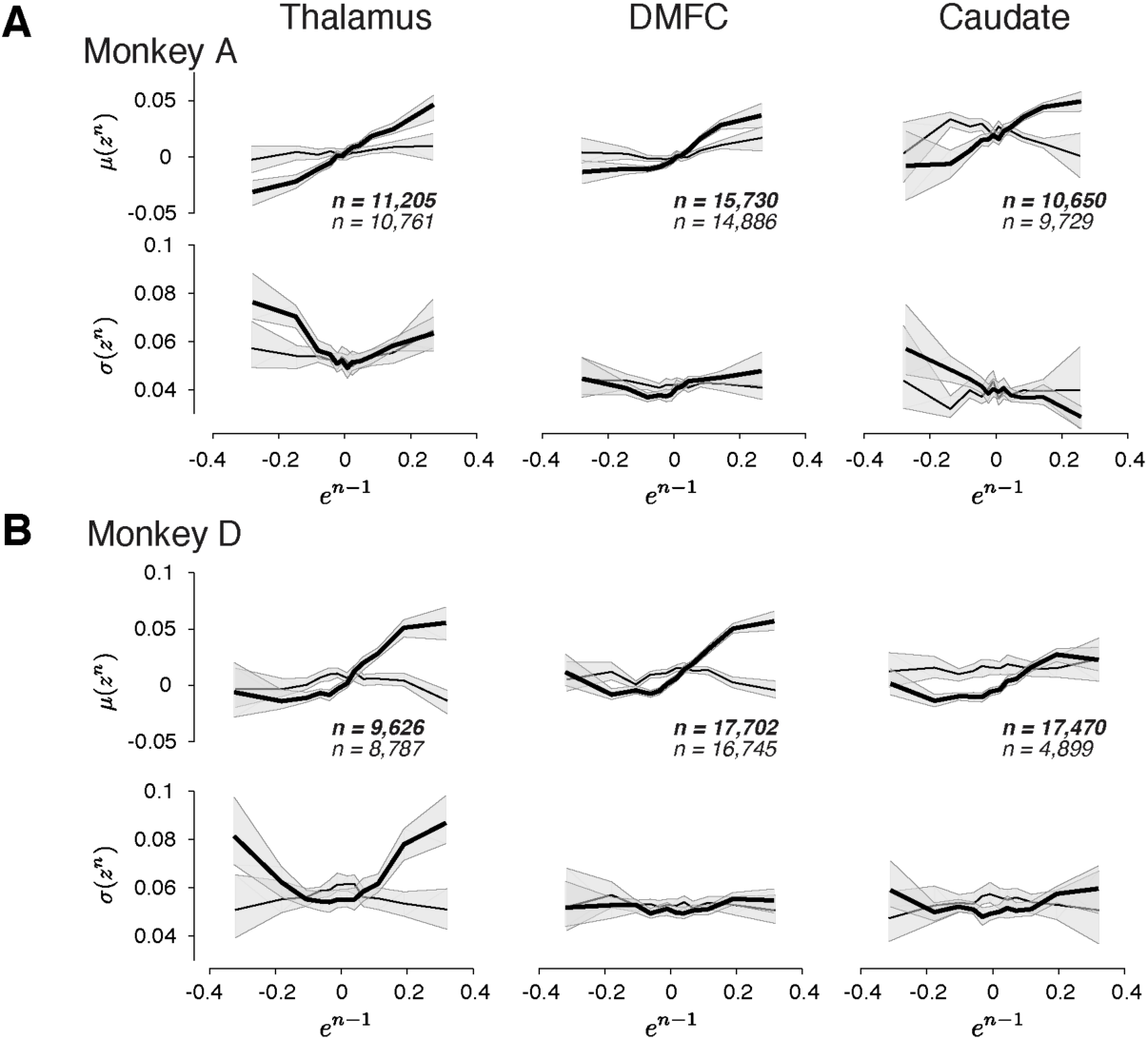
The relationship between neural activity on trial *n* to relative error in the preceding trial (*e*^*n-1*^) in three brain areas and two animals. **(A)** and **(B)** Same format as in Figure 7B-D, with the addition of confidence intervals [1%-99%] (shaded region) obtained by randomly sampling a subset of trials in each bin of *e*^*n-1*^.

**Figure S11.**
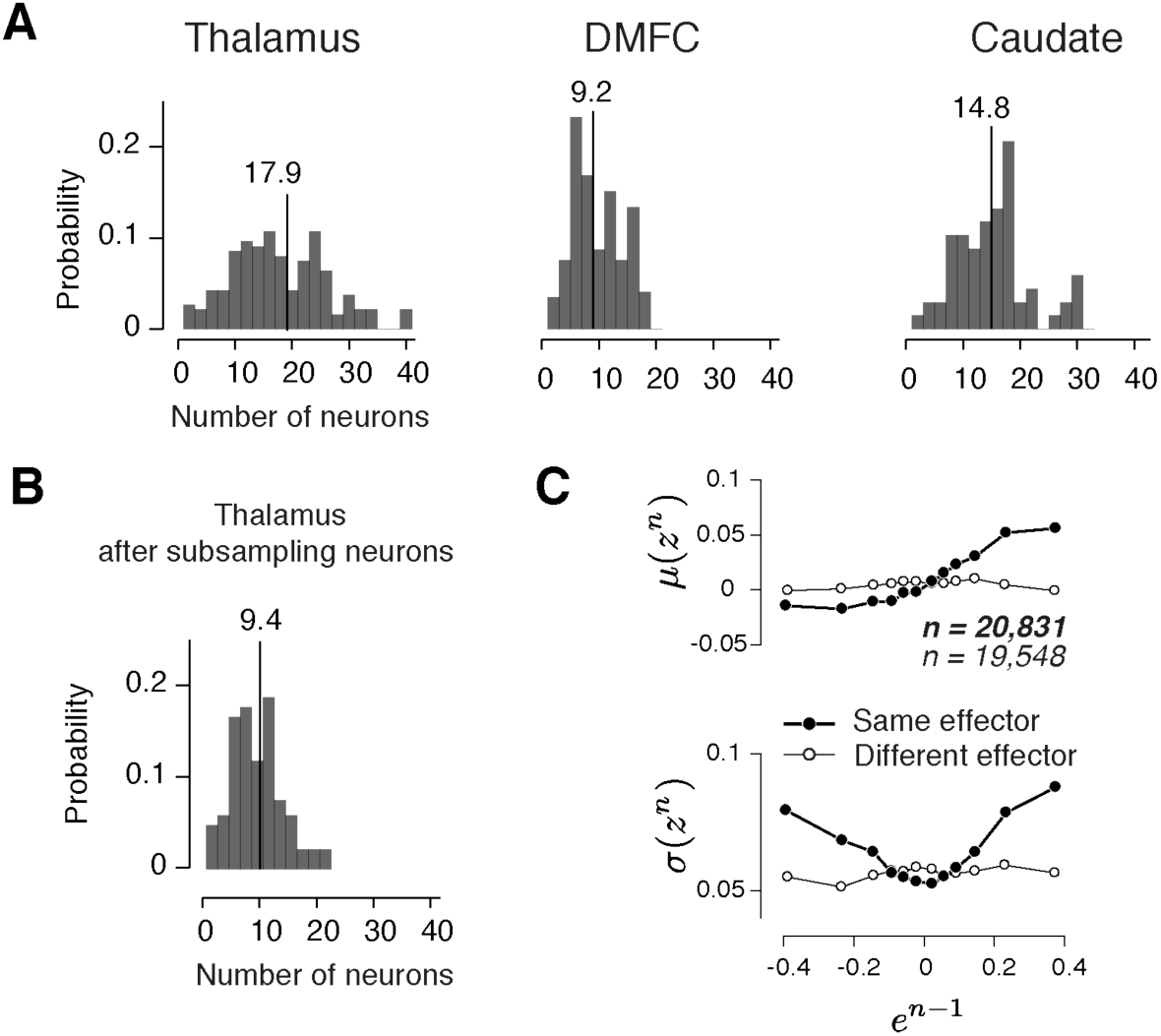
Drift and reward-dependent variability in the three brain areas inferred from comparable number of simultaneously recorded neurons. **(A)** Histogram of the number of simultaneously recorded neurons in the thalamus, DMFC and caudate across sessions. The averages are shown on top. **(B)** The number of simultaneously recorded neurons in the thalamus subsampled by a factor of 2 to approximately match DMFC. **(C)** Analysis of drift and reward-dependent variability for the sub-sampled thalamic population, shown in the same format as Figure 7B.

**Figure S12.**
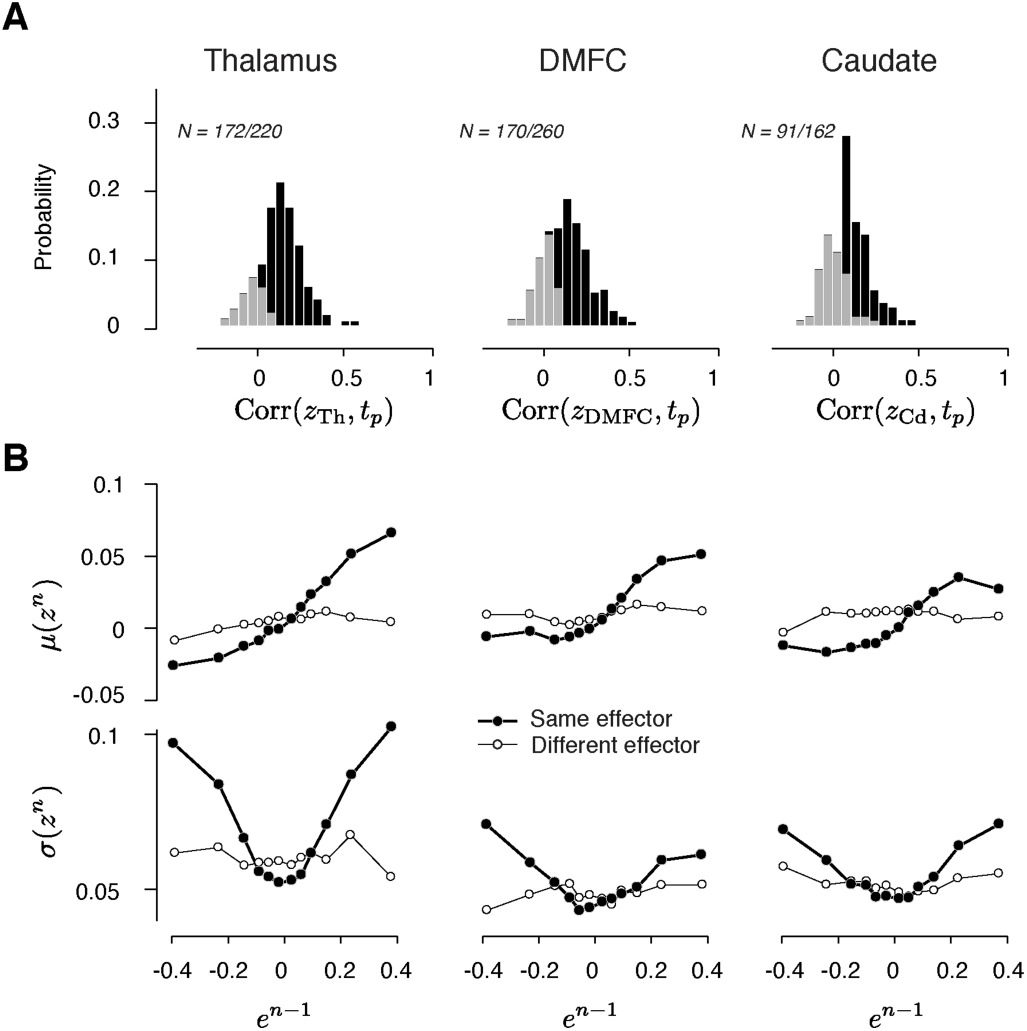
Drift and variability of spike count along the direction that decodes the produced interval (*t*_*p*_) in the thalamus, DMFC and caudate. **(A)** Histogram of the correlation coefficients between *t*_*p*_ and the output of a decoder of *t*_*p*_ from neural data, denoted *z* (black: significantly positive). Results are shown with the same format as in Figure 6B. **(B)** Mean (top; *μ(z*^*n*^)) and standard deviation (bottom; *σ(z*^*n*^)) of projected neural activity as a function of error in the preceding trial (*e*^*n-1*^). Results are shown with the same format as in Figure 7B (thick lines: same effector; thin lines: different effectors).

**Figure S13.**
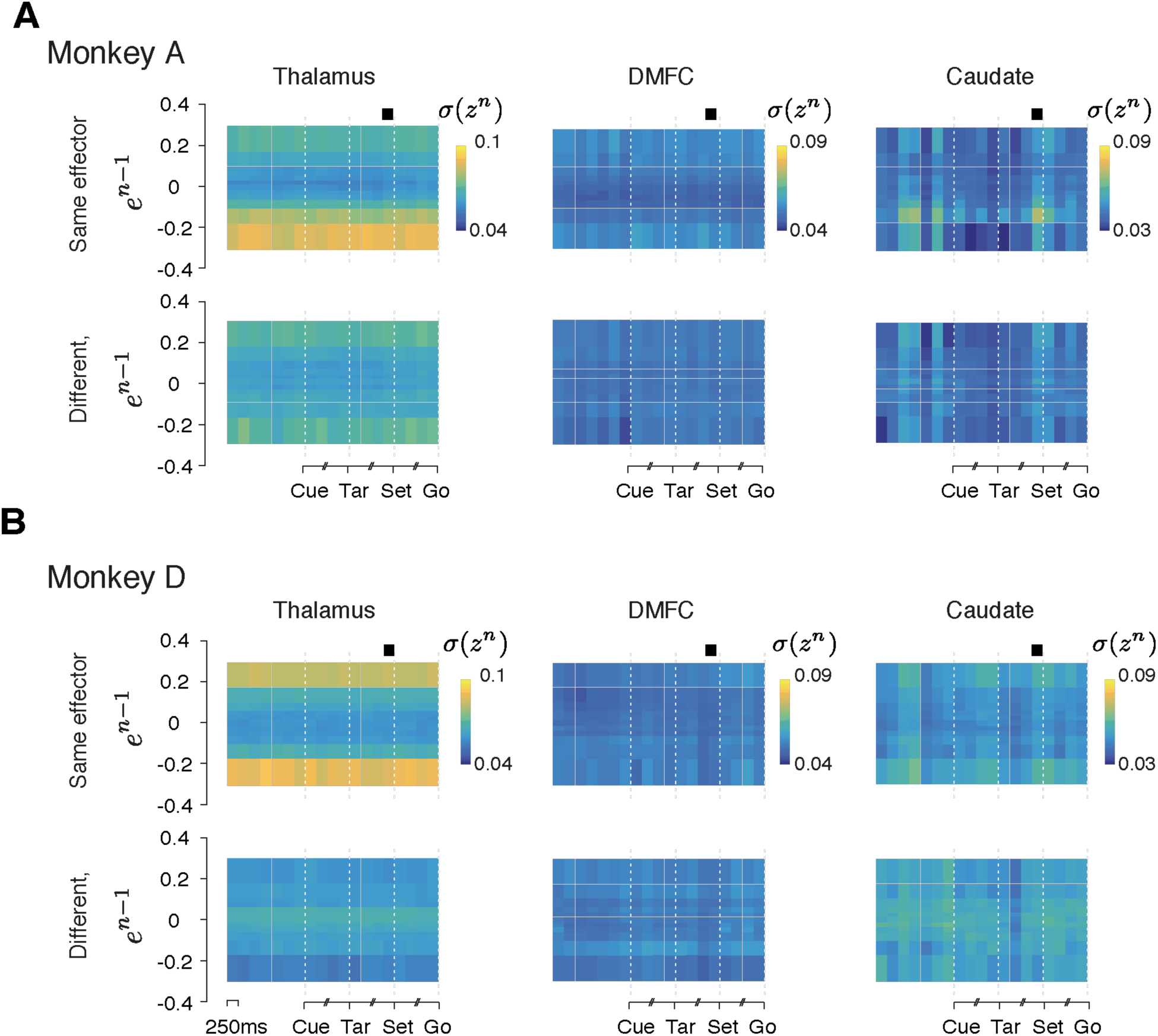
Reward-dependent neural variability throughout the trial. Same as Figure 8 shown for two animals separately.

**Figure S14.**
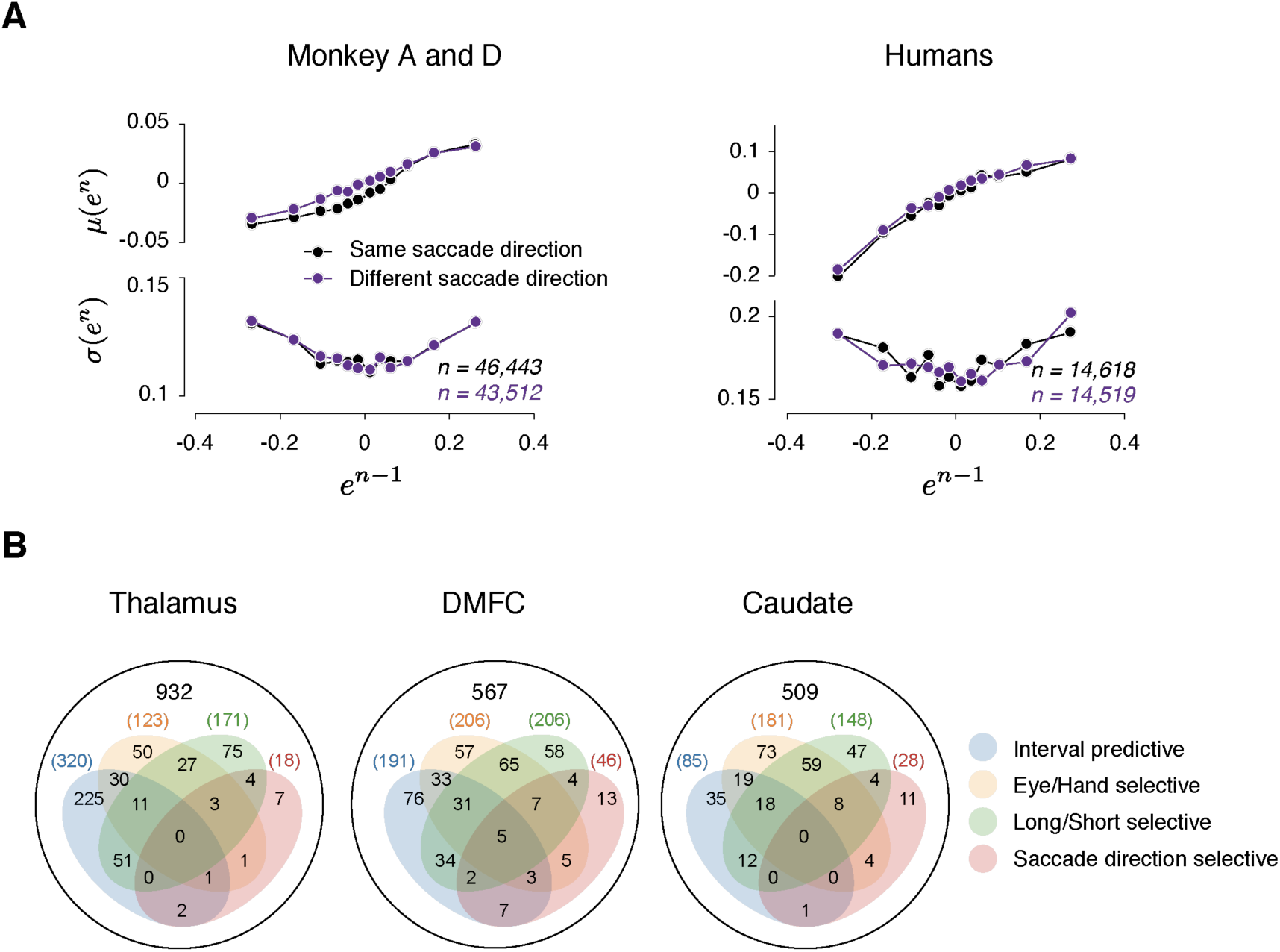
Analysis of neural data in relation to saccade direction. **(A)** Error statistics across consecutive trials. *μ(e*^*n*^) (top) and *σ(e*^*n*^) (bottom) as a function of *e*^*n-1*^. Similar to the main paper, we tested the U-shaped profile using quadratic regression. The coefficients of the squared term was not significantly different between the two directions (Monkeys: 0.26 [0.17, 0.34] vs 0.27 [0.2, 0.34]; Humans: 0.33 [0.063, 0.59] vs 0.4 [0.24, 0.55]). **(B)** A Venn diagram showing the number of individual neurons in three brain areas whose spiking during a 250 ms time window before Set was significantly different across effectors (Eye/Hand; p < 0.01, two-sample *t*-test), target intervals (Short/Long; p < 0.01, two-sample *t*-test), saccade directions (Left/Right; p < 0.01, two-sample *t*-test), and predictive of the produced interval (p < 0.01, shuffling test). The number of neurons in each category is indicated in the diagram. Among the neurons that were predictive of the produced interval (intersection of blue and dark red), 3/320 in the thalamus, 17/191 in DMFC, and 1 out of 85 in the caudate were direction selective.

**Figure S15.**
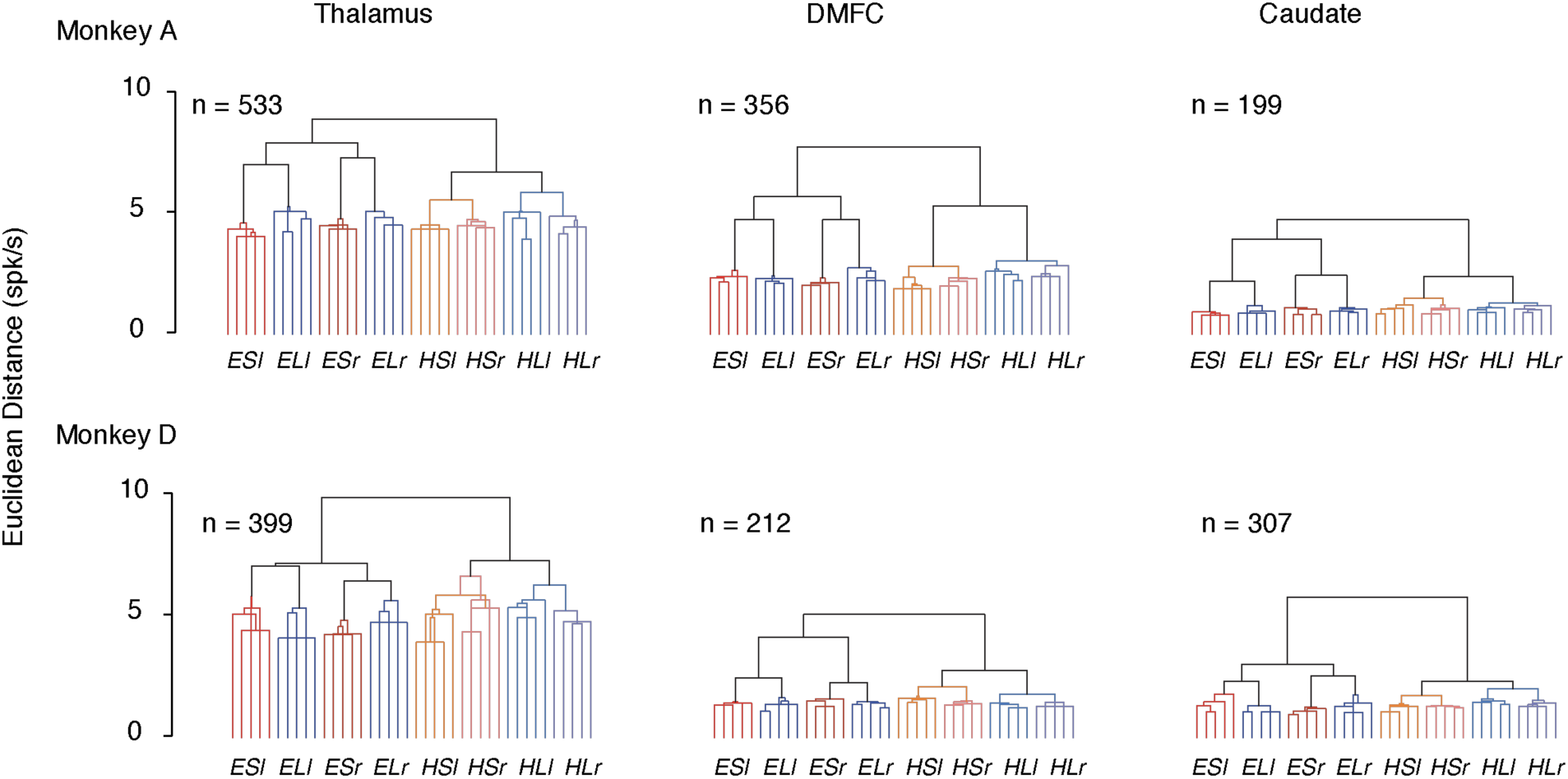
Euclidean distance between spike counts across the population associated with different trial types. Each dendrogram summarizes the Euclidean distance of population spike count between different trial types as shown below the abscissa (E/H: Eye/Hand effector, S/L: Short/Long interval, l/r: left/right saccade direction). Spike counts were obtained from a 250 ms time window before Set in each area (columns, the total number of neurons used were indicated) and each animal (rows). We assessed the reliability of the organization of the dendrogram by applying the same analysis 100 times to 20% of trials randomly sampled from a specific trial type. The structure of the top nodes associated with effector and interval were more consistent across repeats than expected by chance (p<0.01). The plot shows the results for 5 such random repeats. Distances were normalized by the number of neurons in each area and animal (i.e. divided by 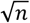 to obtain the average distance per neuron in the state space).

**Figure S16.**
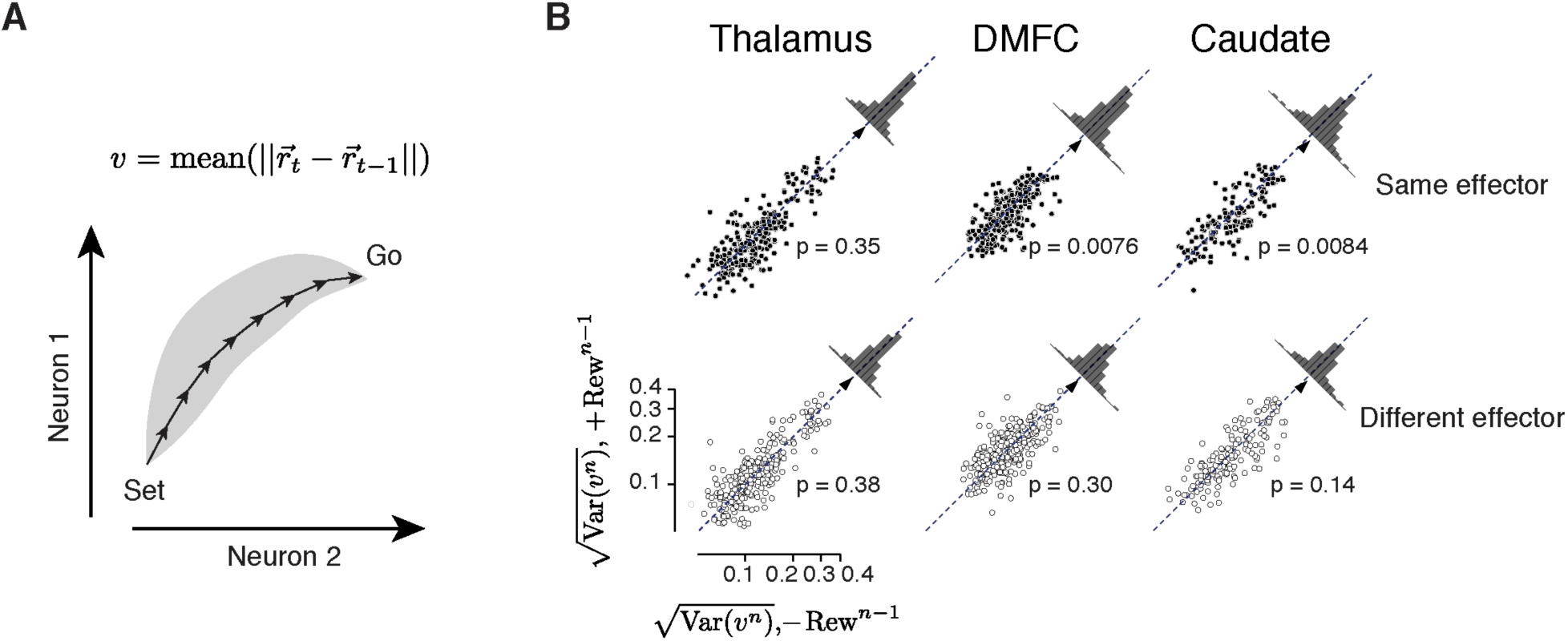
Analysis of speed variability across single trials as a function of reward. **(A)** Schematics showing how we estimated speed of population neural trajectories for single trial (*v*^n^). We estimated the average speed for each trial by averaging the Euclidean distance of population spike counts between nearby points (*dt* = 125 ms) along the trajectory. To minimize the differential contribution of neurons with different average firing rates to speed estimates, spike counts for each neuron was normalized (z-score) across trials and time bins. **(B)** Variability of speed estimates 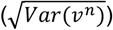 after rewarded trials (+Rew^n-1^, ordinate) compared to unrewarded trials (-Rew^n-1^, abscissa) for consecutive trials of the same (top row) and different effector (bottom row) in the thalamus (left), DMFC (middle) and caudate (right). In every session, each trial type contributed two points to each panel, one for when the preceding error was positive and one for when it was negative. We treated these two error conditions independently so as to minimize any potential dependence of our estimated 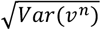 on average speed *v*^n^ that may change depending on the sign of error (Wang *et al.*, 2018). The scatter plot in each panel shows the distribution of speed variability for trials that succeeded a rewarded trial (ordinate) relative to the distribution of speed variability for trials that succeeded an unrewarded trial (abscissa). To test whether the speed was significantly different between these two conditions, we used a two-tail paired sample t-test to determine whether and in which direction the data deviated significantly from the unity line (diagonal). The distribution of the difference between the two conditions is shown on the top right of each panel, and the mean for each distribution is indicated (triangle). P-values for each test are indicated. (Thalamus: same effector, t_271_ = 0.94, different effector, t_271_= −0.88, DMFC: same effector, t_313_ = 2.69, different effector, t_313_ = 1.03, Caudate: same effector, t_161_ = 2.67, different effector, t_161_ = 1.47).

**Table S1.** Ground truth versus model fits for the hyperparameters used in RSGP simulation in Figure 5A and S8A.

| | $l_{SE}$ | $\sigma_{SE}$ | $l_{RS}$ | $\sigma_{RS}$ | $\sigma_0$ |
| --- | --- | --- | --- | --- | --- |
| Ground truth | 20.0 | 0.141 | 2.0 | 0.141 | 0.10 |
| MML fit | 18.4 | 0.129 | 2.14 | 0.137 | 0.0749 |

**Table S2.**
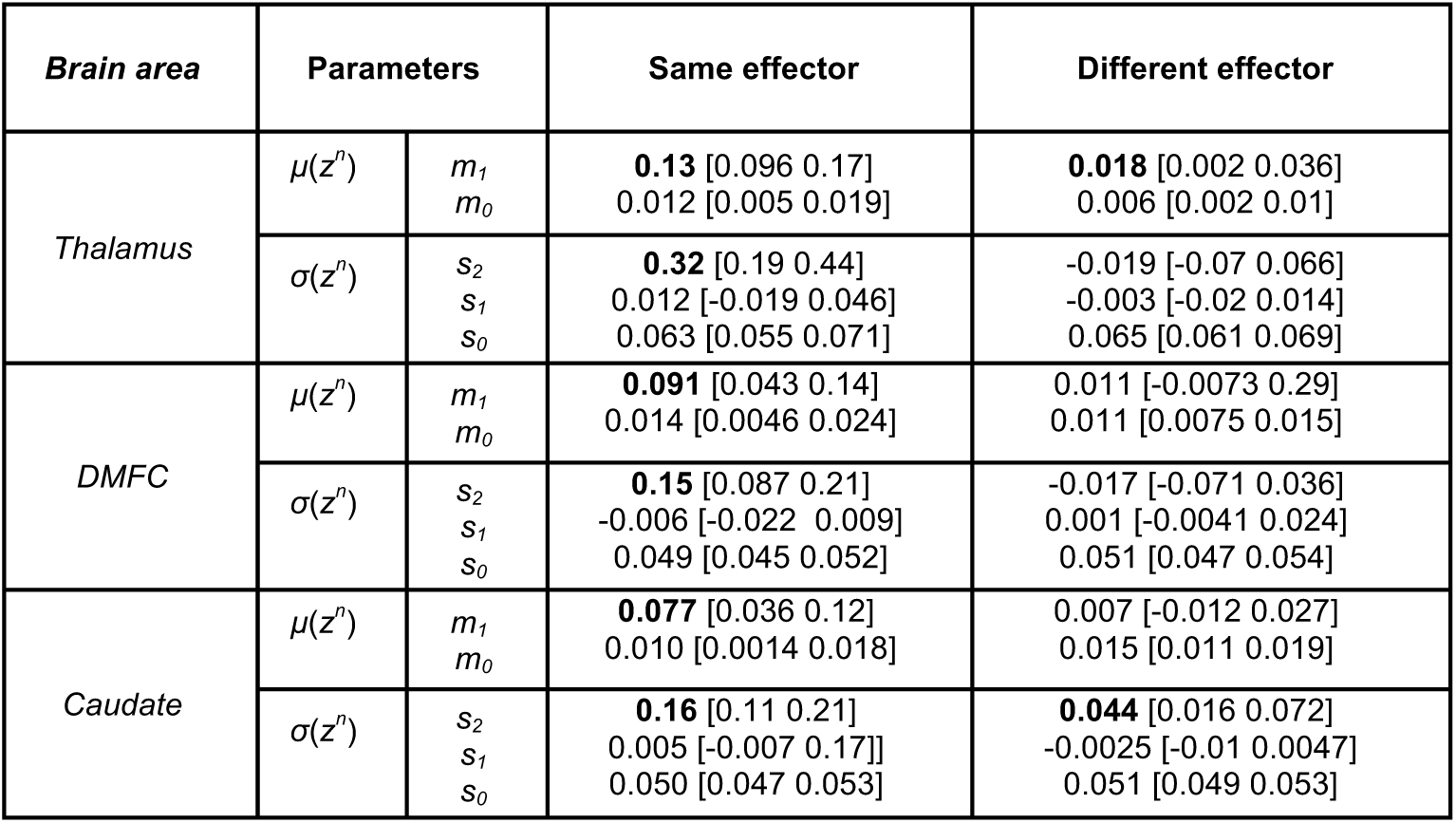
Regression model fits relating spike count along the direction that predicts produced interval (*t*_*p*_) on trial *n* (*z*^*n*^) to error in trial *n-1* (*e*^*n-1*^). *m*_*0*_ and *m*_*1*_ are parameters of the linear regression model relating the mean of *z*^*n*^ (*μ*(*z*^*n*^)) to *e*^*n-1*^; i.e., *μ*(*z*^*n*^) *= m*_*0*_*+m*_*1*_*e*^*n-1*^. *s*_*0*_, *s*_*1*_ and *s*_*2*_ are parameters of the quadratic regression model relating the standard deviation of *z*^*n*^ (*σ*(*z*^*n*^)) to *e*^*n-1*^; i.e., *σ*(*z*^*n*^) *= s*_*0*_*+s*_*1*_*e*^*n-1*^*+s*_*2*_(*e*^*n-1*^)^*2*^. Fit parameters are shown separately for the thalamus, DMFC and caudate and further separated depending on whether trial *n-1* and *n* were of the same or different effectors. Bold indicated significantly positive value (p < 0.01, 1% and 99% confidence intervals of the estimation were shown).

